# Nuclear Translocation of SIRT4 Mediates Deacetylation of U2AF2 to Modulate Renal Fibrosis Through Alternative Splicing-mediated Upregulation of CCN2

**DOI:** 10.1101/2024.04.30.591939

**Authors:** Guangyan Yang, Jiaqing Xiang, Xiaoxiao Yang, Xiaomai Liu, Yanchun Li, Lixing Li, Lin Kang, Zhen Liang, Shu Yang

## Abstract

TGF-β stimulates CCN2 expression which in turn amplifies TGF-β signaling. This process promotes extracellular matrix production and accelerates the pathological progression of fibrotic diseases. Alternative splicing plays an important role in multiple disease development, while U2 small nuclear RNA auxiliary factor 2 (U2AF2) is an essential factor in the early steps of pre-mRNA splicing. However, the molecular mechanism underlying abnormal *CCN2* expression upon TGF-β stimulation remains unclear. This study elucidates that SIRT4 acts as a master regulator for CCN2 expression in response to TGF-β by modulating U2AF2-mediated alternative splicing. Analyses of renal biopsy specimens from patients with CKD and mouse fibrotic kidney tissues revealed marked nuclear accumulation of SIRT4. The tubulointerstitial fibrosis was alleviated by global deletion or tubular epithelial cell (TEC)-specific knockout of *Sirt4*, and aggravated by adeno-associated virus-mediated SIRT4 overexpression in TECs. Furthermore, SIRT4 was found to translocate from the mitochondria to the cytoplasm through the BAX/BAK pore under TGF-β stimulation. In the cytoplasm, TGF-β activated the ERK pathway and induced the phosphorylation of SIRT4 at Ser36, which further promoted its interaction with importin α1 and subsequent nuclear translocation. In the nucleus, SIRT4 was found to deacetylate U2AF2 at K413, facilitating the splicing of CCN2 pre-mRNA to promote CCN2 protein expression. Importantly, exosomes containing anti-SIRT4 antibodies were found to effectively mitigate the UUO-induced kidney fibrosis in mice. Collectively, these findings indicated that SIRT4 plays a role in kidney fibrosis by regulating CCN2 expression via the pre-mRNA splicing.

## 1. Introduction

In the kidney and other organ systems, the overexpression (OE) of cellular communication network 2 (CCN2), also known as connective tissue growth factor, is widely recognized as a marker of fibrotic activity^1^. Transforming growth factor-β (TGF-β) is a master regulator of tissue growth, regeneration, remodeling, and fibrosis, Most TGF-β responses involve CCN2 stimulation at some level, such as the stimulation of extracellular matrix (ECM) components and fibrosis^2,3^. Numerous signaling molecules are involved in the crosstalk and integration of TGF-β and CCN2 effects and vary depending on the cell type and the physiological or pathological process involved. For instance, TGF-β-stimulated SMADs are necessary for the induction of CCN2 expression in normal fibroblasts and basal CCN2 induction in scleroderma fibroblasts, while the maintenance of CCN2 expression is independent of SMADs^4^. However, it remains unknown whether TGF-β1 regulates CCN2 expression via non-transcriptional pathways.

Sirtuins (Sirts), mammalian Sir2 orthologs, are a highly conserved family of nicotinamide adenine dinucleotide-dependent protein deacetylases and act as important regulators of the aging process, inflammation, cancer, and metabolic diseases^5–7^. Among the seven known mammalian Sirts, SIRT4 possesses ADP-ribosyltransferase, lipoamidase, and deacylase activities^8,9^. The inhibition of SIRT4 expression has been shown to increase the fat-oxidation capacity of the liver and mitochondrial function in the muscle^10^. As loss of fatty acid β-oxidation in the proximal tubule is a critical mediator of acute kidney injury and eventual fibrosis^11,12^, we hypothesize that SIRT4 may act as a pro-fibrotic factor. However, other studies have demonstrated that SIRT4 OE inhibits glutamine metabolism^13^, which is necessary for collagen protein synthesis^14^. This suggests a potential protective role of SIRT4 in the context of fibrosis. However, the specific mechanism by which SIRT4 regulates renal fibrosis remains unclear. Therefore, research is needed to further explore the role and mechanisms of SIRT4 in renal fibrosis, as well as potential therapeutic strategies.

An estimated 60% of all human genes undergo alternative splicing, a highly regulated process that produces splice variants with different functions^15^. There are five small nuclear ribonucleoproteins (snRNPs)—U1, U2, U5, and U4/U6. The status of these proteins helps in maintaining sufficient proteomic diversity for the functional requirements of cell fates and body homeostasis^16^. U2 small nuclear RNA auxiliary factor 1 (U2AF1), together with U2AF2, forms the U2AF complex that recognizes and binds to the 3′ splice site of pre-mRNA and recruits U2 snRNPs, thereby facilitating the assembly of the spliceosome, a large RNA-protein complex responsible for the splicing of pre-mRNA^17–20^. The SF3B complex, which is the core of the U2 snRNP, comprises SF3B1, SF3B2, SF3B3, SF3B4, SF3B5, SF3B6, and PHF5A/ SF3B14b^21^. Dysregulation of U2AF2 can lead to the disruption of splicing events, which may contribute to the development and progression of kidney fibrosis. Accordingly, understanding the role of U2AF2 in kidney fibrosis may provide valuable insights into its underlying molecular mechanisms.

Here, we demonstrated that TGF-β1 induced SIRT4 nuclear translocation, resulting in the deacetylation of U2AF2 and recruitment of U2 snRNP, subsequently contributing to pre-mRNA splicing and enhanced protein expression of CCN2. Overall, our study revealed that SIRT4 inhibition can alleviate the progression of renal fibrosis by suppressing CCN2 expression.

## 2. Methods

### 2.1 Studies in animals

All animal care and experimental protocols for in vivo studies conformed to the Guide for the Care and Use of Laboratory Animals published by the National Institutes of Health (NIH; NIH publication no.:85–23, revised 1996). The sample size for the animal studies was calculated based on a survey of data from published research or preliminary studies. *Sirt4^flox/flox^* (C57BL/6J-*Sirt4^em1flox^/*Cya; Strain ID: CKOCMP-75387-*Sirt4*-B6J-VA), Col1a2-Cre/ERT2 mice (Cat#: C001248) and Cdh16-Cre mice (Cat#: C001452) obtained from Cyagen Biosciences (Guangzhou) Inc (Guangzhou, Guangdong, China). C57BL/6J mice and *Sirt4^−/−^* (C57BL/6JGpt-*Sirt4^em13Cd1976^*/Gpt; Strain ID: T011568) mice based on C57BL/6J background were purchased from Gempharmatech Co. Ltd (Jiangsu, Nanjing, China). *U2af2^flox/flox^* (C57BL/6J-*U2af2^em1(flox)Smoc^*) mice based on C57BL/6J background were purchased from Shanghai Model Organisms. These mice were maintained in SPF units of the Animal Center of Shenzhen People’s Hospital with a 12 h light cycle from 8 a.m. to 8 p.m., 23 ± 1 °C, 60 – 70 % humidity. Mice were allowed to acclimatize to their housing environment for 7 days before the experiments. At the end of the experiments, all mice were anesthetized and euthanized in a CO_2_ chamber, followed by the collection of kidney tissues. All animals were randomized before treatment. Mice were treated in a blinded fashion as the drugs used for treating animals were prepared by researchers who did not carry out the treatments. No mice were excluded from the statistical analysis. Studies were performed in accordance with the German Animal Welfare Act and reporting follows the ARRIVE guidelines.

### 2.2 Generation of cell type–specific *Sirt4* conditional knockout mice

To generate fibroblast-specific conditional *Sirt4* knockout mouse line, *Sirt4^flox/flox^* mice were bred with Col1a2-Cre/ERT2 transgenic mice. To activate the Cre-ERT system, tamoxifen (80 mg/kg/day, dissolved in olive oil) was injected intraperitoneally for 4 consecutive days 2 weeks before the induction of renal fibrosis in control and S4FKO mice. After UUO, uIRI, or FA, tamoxifen diet was administered until sacrifice in order to ensure the deletion of Sirt4 in newly generated myofibroblasts. To generate TECs-specific conditional Sirt4 knockout mouse line (S4TKO), *Sirt ^flox/flox^* mice were bred with Cdh16-Cre transgenic mice.

### 2.3 Mouse kidneys were transfected with adeno-associated virus vector (AAV)

8-week-old mice received in situ renal injection with AAV9- empty vector (AAV9-*Ctrl*; control group), AAV9-*Ksp*-*Sirt4* (*Sirt4^OE^* group; *Ksp*, tubule specific promoter), AAV9-*Ksp*-wild type U2af2 (*wtU2af2*), and AAV9-*Ksp*-mutant U2af2 (*mU2af2*) at three independent points (10 - 15 μl virus per poin; virus injected dose: 2.5 E + 11 v.g.) in the kidneys of mice (n = 6). Adeno-associated virus type 9 constructs, including GV501 empty vector, *Sirt4*, *wtU2af2*, and *mU2af2* were provided by GeneChem Company (Shanghai, China).

### 2.4 Mice kidney fibrotic models

Male C57BL/6, *Sirt4*^−/−^, *Sirt ^flox/flox^*, and cell type–specific conditional knockout mice (∼8 to 10 weeks old) were subjected to various kidney injury models to induce renal fibrosis. UUO was performed by permanent ligation of the right ureter with 6-0 silk. Ureter-ligated kidneys and contralateral kidneys (CLs), used as nonfibrotic controls, were collected 10 days after surgery. To establish uIRI, left renal pedicles were clamped with microaneurysm clips for 30 minutes followed by reperfusion. During uIRI surgery, mice were placed on a heating pad to maintain body temperature at 37°C. Injured and contralateral kidneys were collected 1 day or 28 days after surgery for analyses. Folic acid (FA)–induced renal fibrosis was conducted by single intraperitoneal injection of 250 mg/kg folic acid (Sigma-Aldrich, 7876) dissolved in 0.3M sodium bicarbonate, and mice were sacrificed 14 days after FA treatment. Mice injected with sodium bicarbonate served as vehicle control.

### 2.5 Study approval

All animal care and experimental protocols for in vivo studies conformed to the Guide for the Care and Use of Laboratory Animals, published by the National Institutes of Health (NIH; NIH publication no.: 85–23, revised 1996), was approved by the Animal Care Committees of the First Affiliated Hospital of Southern University of Science and Technology (No. AUP-230809-LZ-0426-01), and were performed in compliance with the ARRIVE guidelines. Studies with human participants were conducted in line with the Declaration of Helsinki. The studies were approved by Ethics Committee of the First Affiliated Hospital of Southern University of Science and Technology. The written consent obtained from patients was informed consent.

### 2.6 Quantification and Statistical Analysis

All data were generated from at least three independent experiments. Each value was presented as the mean ± SD. All raw data were initially subjected to a normal distribution and analysis by one-sample Kolmogorov-Smirnov (K-S) nonparametric test using SPSS 22.0 software. For animal and cellular experiments, a two-tailed unpaired student’s t-test was performed to compare the two groups. One-way ANOVA followed by the Bonferroni’s post-hoc test was used to compare more than two groups. To avoid bias, all statistical analyses were performed blindly. Statistical significance was indicated at \**P* < 0.05, \*\**P* < 0.01, and \*\*\**P* < 0.001.

## 3. Results

### 3.1. Nuclear localization of SIRT4 increases in fibrotic kidney

The application of mitochondrial SIRT4 tripartite abundance reporter, a tripartite probe for visualizing the distribution of SIRT4 between mitochondria and the nucleus in single cells^22^, proved the importation of SIRT4 into the mitochondrial matrix and demonstrated its localization in the nucleus under mitochondrial stress conditions. Nuclear accumulation of SIRT4 was observed in the kidneys following unilateral ureteral occlusion (UUO) or unilateral renal ischemia-reperfusion injury (uIRI) surgery **(Fig. 1A–D)**. Consistently, elevated nuclear accumulation of SIRT4 was observed in kidney sections from patients with chronic kidney disease (CKD) with severe collagen deposition **(Fig. 1E, F)**.

**Figure 1.**
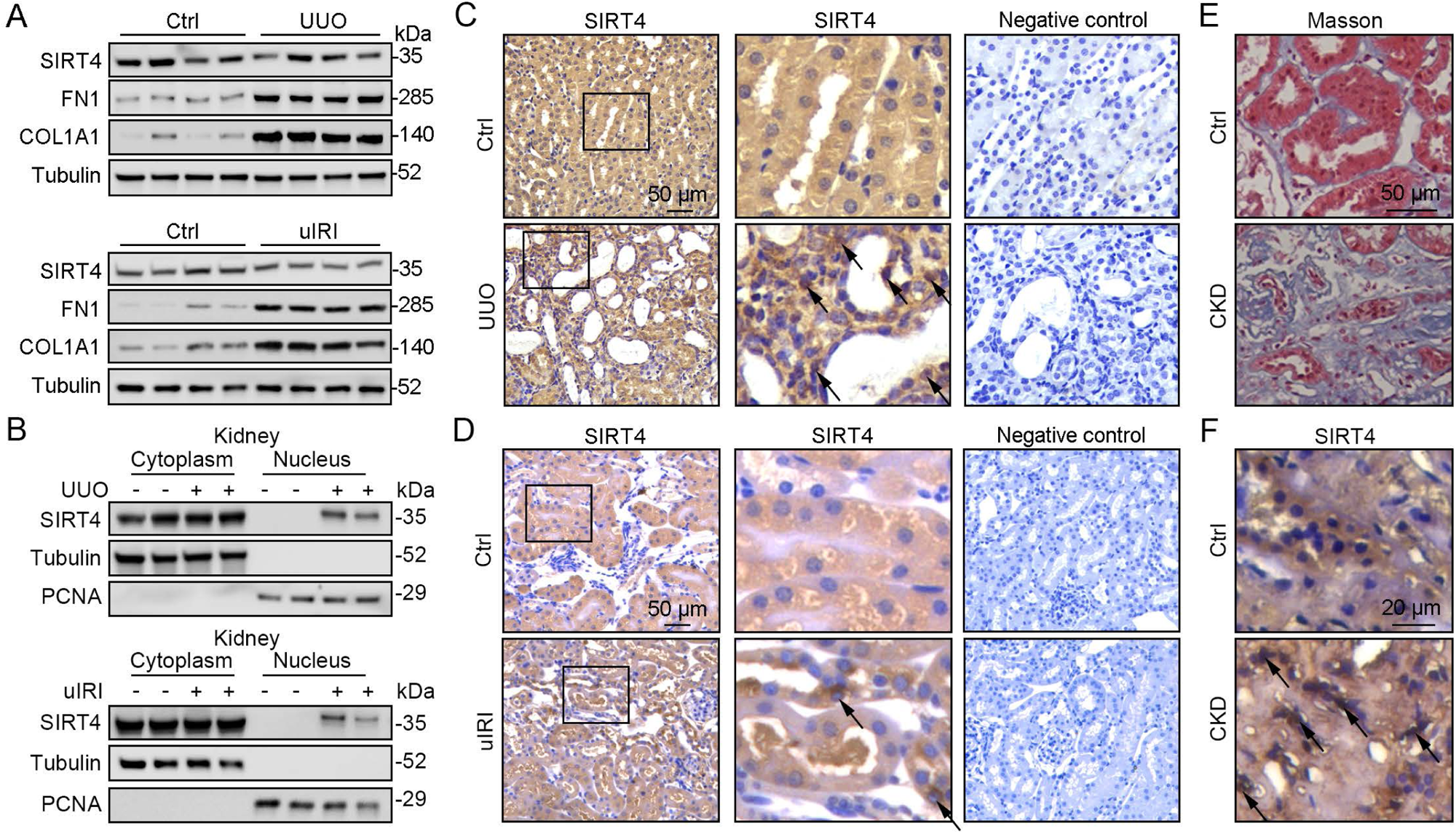
Nuclear accumulation of SIRT4 is increased in renal tubules after injury. **(A)** Western blots analysis of SIRT4, FN1, COL1A1, and Tubulin in the kidney of UUO, uIRI, and sham mice (control group). **(B)** Nuclear fractions were prepared from the kidney of UUO, uIRI, and sham mice. Nuclear PCNA and cytoplasmic tubulin were used as controls. **(C, D)** Representative images of immunohistochemical staining of SIRT4 (scale bar, 50 μm) in the kidneys from mice that underwent sham surgery, UUO surgery on day 10 post surgery or uIRI surgery on day 28 post surgery. **(E, F)** Representative images of Masson’s trichrome staining (the upper panel; scale bar = 50 μm) and SIRT4 immumohistochemical staining (the bottom panel; scale bar = 20 μm) in the kidney sections from patients with CKD (n = 8) and minimal change disease (control group, n = 1).

### 3.2 Global deletion of *Sirt4* protects against kidney fibrosis

To determine the role of SIRT4 in kidney fibrosis development *in vivo*, wild-type (WT) and *Sirt4* global knockout (S4KO) mice were subjected to UUO, uIRI, and folic acid (FA) treatment. Sirius red and Masson’s trichrome staining of the kidney sections revealed extensive renal fibrosis in WT mice following UUO **(Fig. 2A)**, uIRI **(Fig. 2E)**, and FA administration **(Fig. S1B)**. In contrast, the global deletion of *Sirt4* resulted in a remarkable reduction in the extent of renal fibrosis in all three kidney injury models **(Fig. 2A, E and S1B)**. Immunoblots of whole-kidney tissue lysates showed that UUO, uIRI, and FA treatment led to induced levels of the markers of kidney fibrosis (CCN2, FN1, COL1A1, COL3A1, and α-SMA), while the expression of E-cadherin, a hallmark of epithelial-mesenchymal transition, was decreased, with the changes being more pronounced in WT mice than in S4KO mice **(Fig. 2B, F and S1A)**. Consistent with the attenuated post-injury fibrotic response in S4KO mouse kidneys observed via imaging studies, deletion of *Sirt4* also mitigated the upregulation of the profibrotic genes *Col1a1*, *Fn1*, *Eln*, *Ccn2*, *Acta2*, and *Col3a1* **(Fig. 2C, G and S1C)**. In addition, the mRNA levels of both neutrophil gelatinase-associated lipocalin (*Ngal*) and kidney injury molecule 1 (*Kim-1*), which are markers of acute kidney injury, were significantly increased in the kidney tissues from WT and S4KO mice following injury **(Fig. 2D, H and S1D)**. Notably, the levels of *Ngal* and *Kim-1* were significantly reduced in S4KO mice compared to those in WT mice following injury **(Fig. 2D, H and S1D)**. Collectively, these results suggest that *Sirt4* deletion protects mice against renal fibrosis.

**Figure 2.**
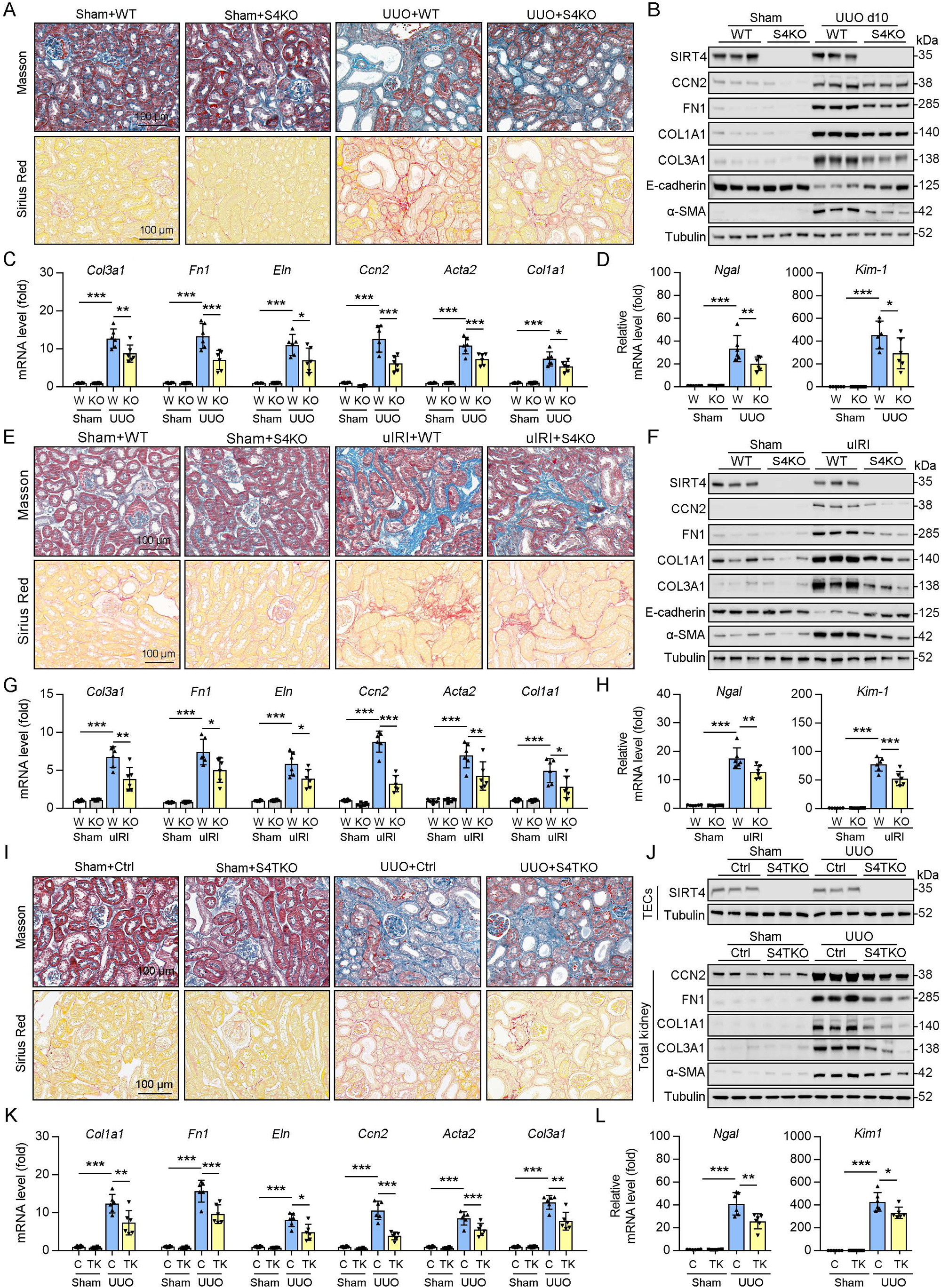
Knockout of *Sirt4* or targeted deletion of *Sirt4* in renal TECs alleviates renal fibrosis induced by UUO or uIRI. **(A to D)** WT or S4KO mice were randomly assigned to sham or UUO surgery according to an established protocol. Kidney samples were obtained from mice on day 10 post UUO or sham surgery (n = 6 per group). A: Representative images of Masson’s trichrome staining and Sirius red in kidneys from mice (scale bar = 100 μm). B: Western blot analysis of the expression of SIRT4, CCN2, FN1, COL1A1, COL3A1, E-cadherin, α-SMA, and Tubulin in the kidney of mice. C, D: The mRNA level of *Col1a1, Fn1, Eln, Ccn2, Acta2*, *Col3a1, Ngal*, and *Kim-1* in the kidney of mice. **(E to H)** WT or S4KO mice received uIRI surgery, and the sham surgery kidneys were used as control. Kidney samples collected from mice after surgery at 28 d (n = 6 per group). E: Representative images of Masson’s trichrome staining and Sirius red in kidneys from mice (scale bar = 100 μm). F: Western blot analysis of the expression of SIRT4, CCN2, FN1, COL1A1, COL3A1, E-cadherin, α-SMA, and Tubulin in the kidney of mice. G, H: The mRNA level of *Col1a1, Fn1, Eln, Ccn2, Acta2*, *Col3a1, Ngal*, and *Kim-1* in the kidney of mice. **(I to L)** WT or S4TKO mice were randomly assigned to sham or UUO surgery according to an established protocol. Kidney samples were obtained from mice on day 10 post UUO or sham surgery (n = 6 per group). **I:** Representative images of Masson’s trichrome staining and Sirius red in kidneys from mice (scale bar = 100 μm). **J:** Western blot analysis of the expression of SIRT4, CCN2, FN1, COL1A1, COL3A1, α-SMA, and Tubulin in the kidney of mice. **K, L:** The mRNA level of *Col1a1, Fn1, Eln, Ccn2, Acta2*, *Col3a1, Ngal*, and *Kim-1* in the kidney of mice. For all panels, data are presented as mean ± SD. \**P* < 0.05, \*\**P* < 0.01, \*\*\**P* < 0.001 by one-way ANOVA with Bonferroni correction test.

### 3.3. Deletion of *Sirt4* in renal tubule epithelial cells markedly attenuates the extent of kidney fibrosis following injury

Using conventional agarose gel-based RT-PCR and Western blot analysis, we analyzed the expression of SIRT4 in renal parenchymal cells, including mouse podocytes (MPCs), glomerular endothelial cells (GECs), kidney fibroblasts (KFs), and tubule epithelial cells (TECs). Compared to their levels in TECs, the basal levels of SIRT4 in MPCs and GECs were relatively low, while in KFs is moderate **(Fig. S1E)**. To determine the *in vivo* contribution of SIRT4 in renal TECs and fibroblasts to the development of kidney fibrosis, we generated TEC-specific (S4TKO; Cdh16-cre/ERT2×*Sirt4^flox/flox^*) and fibroblast-specific (S4FKO; Col1a2-Cre/ERT2×*Sirt4^flox/flox^*) *Sirt4* knockout mice; in these experiments, Cadh16-cre mice and tamoxifen-treated Col1a2-Cre mice were used as controls, respectively. The extent of kidney fibrosis induced by UUO was markedly reduced in S4TKO mice compared to that in control mice **(Fig. 2I, J)**. In addition, the expression levels of kidney fibrosis markers (CCN2, FN1, COL1A1, COL3A1, and α-SMA) were also significantly reduced in S4TKO compared to those in control mice following UUO surgery **(Fig. 2J)**. Consistently, the profibrotic genes (*Col1a1, Fn1, Eln, Ccn2, Acta2*, and *Col3a1*) and acute kidney injury markers *Nagl* and *Kim-1* were downregulated in S4TKO mice compared to those in control mice **(Fig. 2K, L)**. Nevertheless, the targeted deletion of SIRT4 in fibroblasts did not affect the extent of UUO-induced kidney fibrosis **(Fig. S1F–H)**. These results suggest that SIRT4 expressed in TECs but not in fibroblasts primarily contributes to the pathogenesis of kidney fibrosis.

### 3.4 SIRT4 OE aggravates kidney fibrosis

Since the deletion of *Sirt4* attenuated the post-injury fibrotic response, we next determined whether *Sirt4* OE can aggravate the response. We introduced SIRT4 into WT mouse kidneys using the adeno-associated virus serotype 9 vector with a tubule-specific *Ksp-cadherin* promoter, and AAV9-*Ksp*-*null* transfection was used as the control treatment. Targeting *Sirt4* OE in kidney TECs markedly aggravated the extent of kidney fibrosis induced by UUO, uIRI, and FA treatment compared to that in the control mice **(Fig. 3A, D, G)**. The expression levels of kidney fibrosis markers (CCN2, FN1, COL1A1, COL3A1, and α-SMA) and the decreased E-cadherin were also significantly enhanced in *Sirt4* OE mice compared to those in the control mice following kidney injury **(Fig. 3B, E, H)**. Consistently, *Sirt4* OE upregulated the transcription of profibrotic genes, namely *Col1a1*, *Fn1*, *Eln*, *Ccn2, Acta2*, and *Col3a1*, in mouse kidneys following injury, compared to that in the control group **(Fig. 3C, F, I)**. Further analysis revealed an evident upregulation of *Ngal* and *Kim-1* transcripts in the kidneys of *Sirt4* OE mice compared to that in the kidneys of control mice in response to UUO **(Fig. 3C)**. Notably, targeting WT *Sirt4* OE in kidney TECs markedly reversed the decline in collagen deposition and the reduction in ECM-related protein expression in S4KO mice **(Fig. S2A–C)**. These results demonstrate that interventional SIRT4 OE in renal TECs exacerbates kidney fibrosis progression.

**Figure 3.**
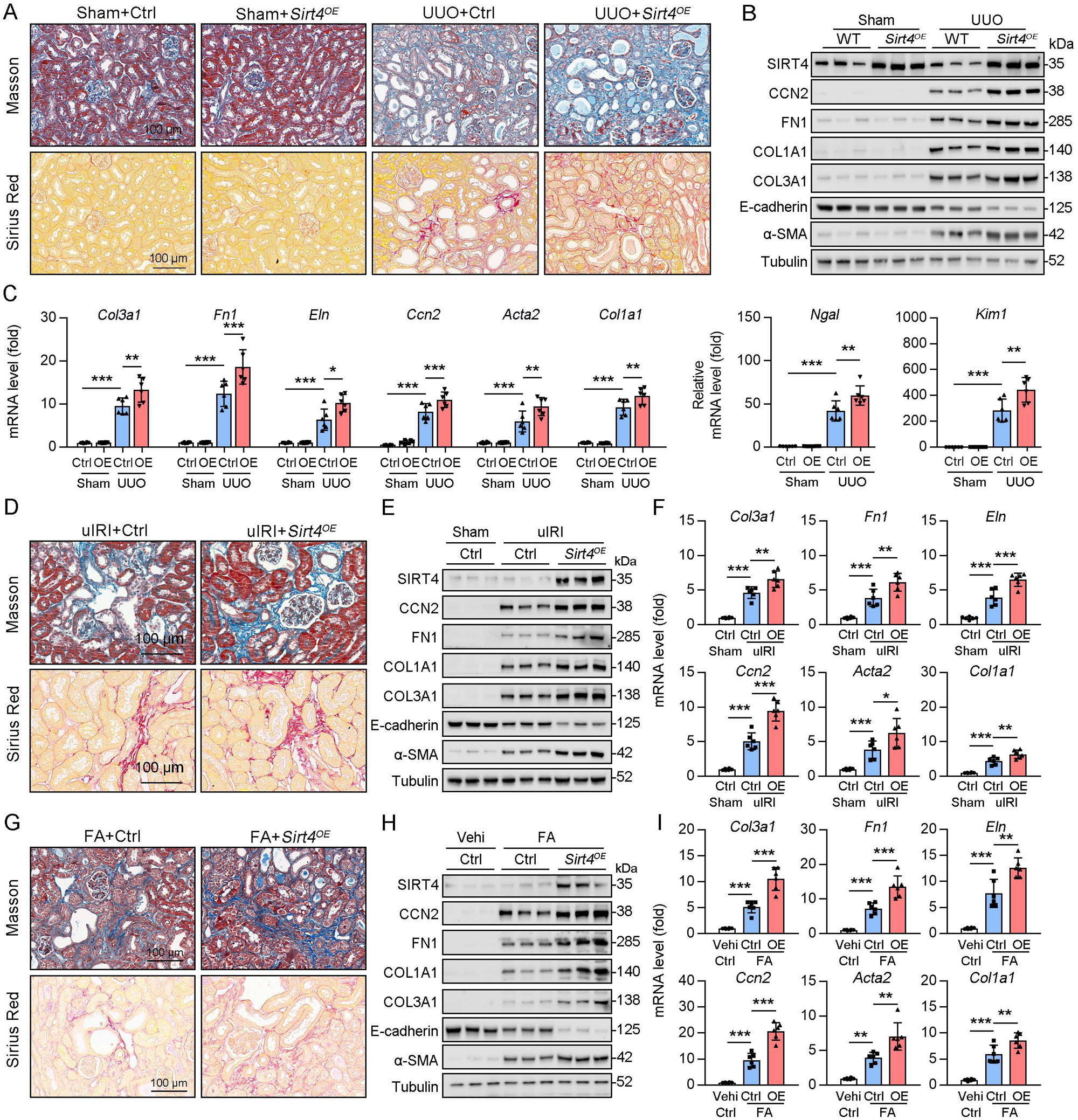
TECs specific SIRT4^OE^ aggravates kidney fibrosis. **(A to I)** AAV9-Ctrl or AAV9-*Ksp*-*Sirt4* was injected into the kidneys of mice in situ at three independent points. After 2-week transfection, the mice received UUO surgery, uIRI surgery, or FA treatment (vehicle treatment as control) (n = 6 per group). A, D, G: Representative images of Masson’s trichrome staining and Sirius red in kidneys from mice (scale bar = 100 μm). B, E, H: Western blot analysis of the expression of SIRT4, CCN2, FN1, COL1A1, COL3A1, E-cadherin, α-SMA, and Tubulin in the kidney of mice. C, F, I: The mRNA level of *Col3a1, Fn1, Eln, Ccn2, Acta2*, *Col1a1, Ngal*, and *Kim-1* in the kidney of mice. For all panels, data are presented as mean ± SD. \**P* < 0.05, \*\**P* < 0.01, \*\*\**P* < 0.001 by one-way ANOVA with Bonferroni correction test.

### 3.5. SIRT4 interacts with U2AF2 under TGF-β stimulation

To explore the action mechanism of SIRT4 in kidney fibrosis, we first identified the proteins interacting with SIRT4 in TECs. We employed rapid immunoprecipitation and mass spectrometry of endogenous proteins (RIME), which is an efficient and unbiased proteomic approach for identifying interacting proteins. Human TECs were treated with TGF-β1 or the control (DMSO) for 24 h, followed by protein-DNA crosslinking in 1% formaldehyde. Cells were sonicated, followed by immunoprecipitation with an SIRT4 antibody **(Fig. 4A)**. Mass spectrometry identified a total of 1081 unique proteins that copurified with SIRT4. We only considered SIRT4-associated proteins that occurred in three of three independent replicates and excluded any proteins that appeared in any one of the IgG control RIMEs. As a result, we selected of 22 SIRT4-associated proteins. Notably, the SIRT4 interaction involved numerous specific nuclear localization proteins, such as PUF60, U2AF2, RPS2, TSR1, ZC3H15, and SART1. Of these, U2AF2 and PUF60 function cooperatively in pre-mRNA splicing^23^. The levels (normalized) of PUF60 and U2AF2 are shown in the **Figure 4B**. We next determined whether PUF60 or U2AF2 was recruited to SIRT4 in TECs stimulated with TGF-β1. Immunoblotting of immunoprecipitated SIRT4 with anti-PUF60 and -U2AF2 antibodies showed that TGF-β1 treatment resulted in an interaction among SIRT4, PUF60, and U2AF2 **(Fig. 4C, left panel)**. Whole-kidney tissue lysates consistently showed an interaction among SIRT4, PUF60, and U2AF2 **(Fig. 4C, middle panel)**. As U2AF2 has been shown to interact with PUF60^23^, we tested whether SIRT4 interacts with either of the two proteins via the U2AF2-PUF60 interaction. The interaction between SIRT4 and PUF60 was abolished in U2af2-knockout cells **(Fig. 4D)**. However, the interaction of SIRT4 and U2AF2 was not affected in *Puf60* knockdown cells (data not shown). Furthermore, exogenous co-IP revealed an interaction between SIRT4 and U2AF2 **(Fig. 4C, right panel)**. Together, these results suggest that SIRT4 interacts with U2AF2 under TGF-β1 stimulation or kidney injury.

**Figure 4.**
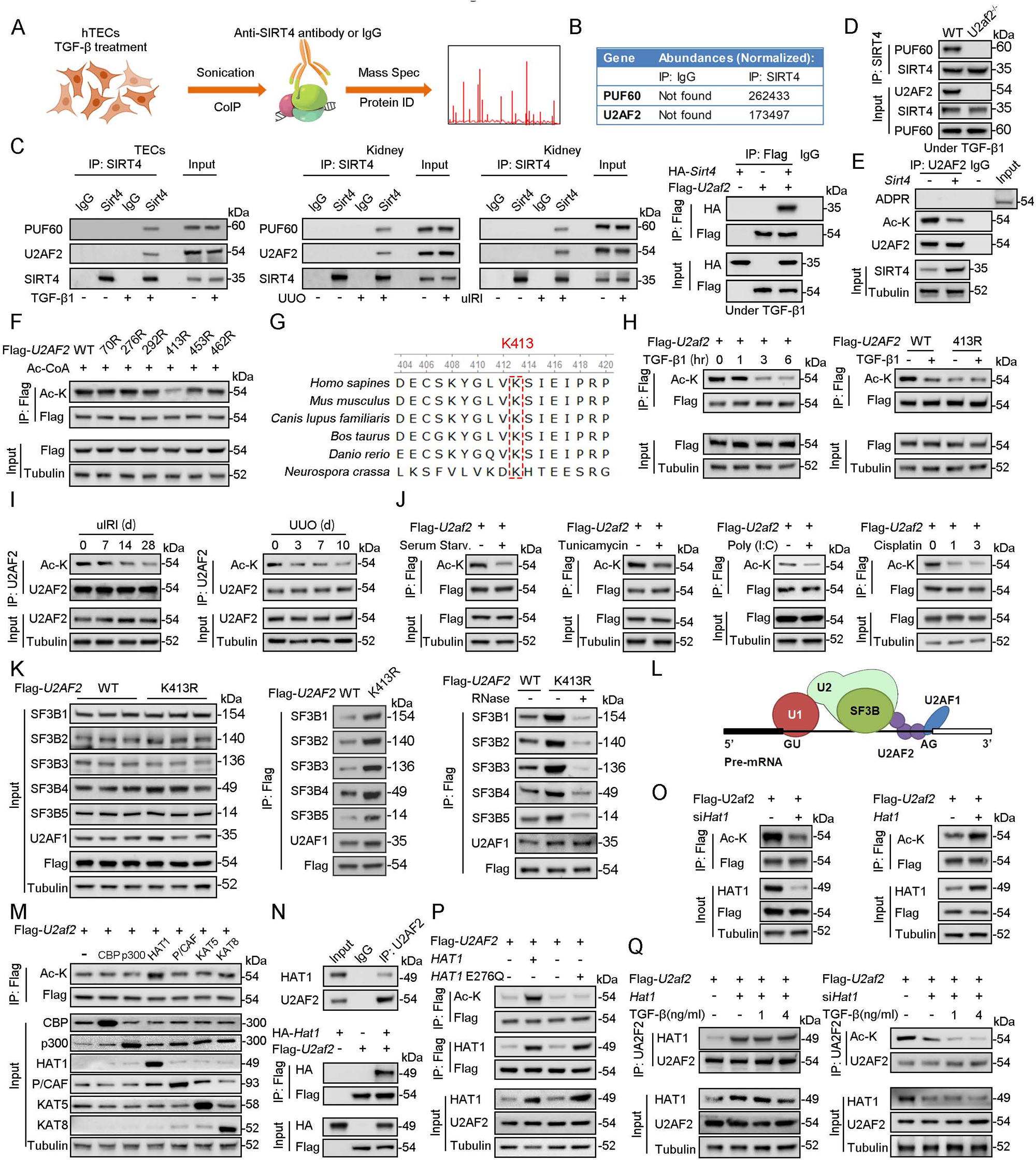
U2AF2 acetylation at K413 is increased under TGF-β1 stimulation. **(A)** Proteins interacted with SIRT4 in chromatin were identified by RIME. **(B)** The abundances (Normalized) of PUF60 and U2AF2 were shown. **(C)** Protein lysates from TECs were exposed to TGF-β1(left panel), kidney from the sham, uIRI, and UUO mice (middle panels), and WT TECs transfected with HA-*Sirt4* and/or Flag-*U2af2* and then treated with TGF-β1 (right panel), the protein lysates were subjected to Co-IP with anti-SIRT4 antibody, and determined protein expression using Western blotting by the indicated antibodies. **(D)** TECs isolated from WT or *U2af2^−/−^*mice and then treated with TGF-β1, cells lysates subjected to Co-IP with anti-SIRT4 antibody, and Western blotting using indicated antibodies. **(E)** WT TECs transfected with Ad-*Sirt4* or Ad-null as indicated, and cells lysates subjected to Co-IP with anti-U2AF2 antibody, and western blotting using indicated antibodies. **(F)** HK2 cells (human renal TECs) transfected with Flag- U2AF2 WT, K70R, K276R, K272R, K292R, K413R, K453R, or K462R under Ac-CoA treatment, cells lysates subjected to Co-IP with anti-Flag antibody, and Western blotting using indicated antibodies. **(G)** The conservation of U2AF2 lysine 413 in different species. **(H)** TECs transfected with Flag-*U2af2* WT under TGF-β1 stimulation, cells lysates subjected to Co-IP with anti-Flag antibody, and western blotting using indicated antibodies (left panel). HK2 cells transfected with Flag-*U2AF2* WT or K413R with or without TGF-β1, cells lysates subjected to Co-IP with anti-Flag antibody, and western blotting using indicated antibodies (right panel). **(I)** Kidney lysates subjected to Co-IP with anti-U2AF2 antibody in uIRI, and UUO mice, and western blotting using indicated antibodies (middle and right panels). **(J)** WT TECs transfected with Flag-*U2af2* WT and then under serum starvation, Tunicamycin, Poly (I:C), and Cisplatin stimulation. Cells lysates subjected to Co-IP with anti-Flag antibody, and Western blotting using indicated antibodies. **(K)** HK2 cells transfected with Flag-*U2AF2* WT or K413R and cells lysates subjected to Co-IP with anti-Flag antibody, and Western blotting using indicated antibodies (left and middle panels). HK2 cells transfected with Flag-*U2AF2* WT or K413R with or without RNase stimulation, cells lysates subjected to Co-IP with anti-Flag antibody, and western blotting using indicated antibodies (right panel). **(L)** Schematic representation of components of SF3B complex performing pre-mRNA BPS recognition. **(M)** WT TECs transfected with Flag-*U2af2* WT with CBP, p300, HAT1, P/CAF, KAT5, or KAT8, cells lysates subjected to Co-IP with anti-Flag antibody, and Western blotting using indicated antibodies. **(N)** Whole-kidney lysates were immunoprecipitated with anti-U2AF2 antibody, and precipitated proteins were detected by indicated antibodies (the upper panel). WT TECs was transfected with HA-*Hat1* and/or Flag-*U2af2*. Cells lysates subjected to Co-IP with anti-Flag antibody, and Western blotting using indicated antibodies. **(O)** WT TECs was transfected with Flag-*U2af2* WT with Ad-sh*Hat1* or Ad-*Hat1*. Cells lysates subjected to Co-IP with anti-Flag antibody, and Western blotting using indicated antibodies. **(P)** HK2 cells were transfected with Flag-*U2AF2* WT, Ad-*HAT1*, or Ad-*HAT1* E276Q as indicated in figure. Cells lysates subjected to Co-IP with anti-Flag antibody, and western blotting using indicated antibodies. **(Q)** WT TECs was transfected with Flag-*U2af2*, Ad-*Hat1* or Ad-sh*Hat1* under TGF-β1 stimulation. Cells lysates subjected to Co-IP with anti-Flag antibody, and western blotting using indicated antibodies.

### 3.6. U2AF2 acetylation is decreased under cellular stress

Accumulating evidence suggests that SIRT4 can show weak ADP-ribosyltransferase^8,24,25^ as well as substrate-specific deacetylase activity^26–28^, akin to that observed for SIRT6 and SIRT7^29,30^. Hence, we tested whether SIRT4 regulates the ADP-ribosylation or acetylation of U2AF2. Our results showed that *Sirt4* OE reduced the levels of acetylated U2AF2 **(Fig. 4E)**. However, the ADP-ribosylation of U2AF2 was extremely low in both SIRT4 OE and control cells **(Fig. 4E)**. According to the database Compendium of Protein Lysine Modifications 4.0, lysine (K) 70, 276, 292, 413, 453, and 462 on U2AF2 can be acetylated. Hence, we mutated all six K residues to arginine (R), which mimics the deacetylated state of the protein. As shown in **Figure 4F**, the U2AF2-K413R mutant showed reduced acetylation compared to U2AF2 WT. Conservation analysis of U2AF2 indicated that K413 is a highly conserved site spanning from *Schizosaccharomyces pombe* to *Homo sapiens* **(Fig. 4G)**. Next, we examined whether U2AF2 K413 is the key acetylation site in response to TGF-β1 stimulation. TGF-β1 treatment reduced the levels of acetylated U2AF2 in WT but had little effect in the K413R mutant U2AF2 **(Fig. 4H)**. Consistently, U2AF2 acetylation decreased in the kidney following injury **(Fig. 4I)**. Interestingly, U2AF2 acetylation was also remarkably reduced in the cells subjected to serum starvation, endoplasmic reticulum (ER) stress (induced by tunicamycin), viral infection [poly (I:C)], or DNA damage stress (induced by cisplatin) **(Fig. 4J)**. These results collectively demonstrate that U2AF2 deacetylation occurs universally in response to cellular stress.

### 3.7. U2AF2 acetylation affects the interaction of U2AF and SF3B complexes

Next, we investigate the effect of U2AF2 acetylation at K413 on the formation of the U2AF complex and recruitment of the SF3B complex **(Fig. 4K)**. The expression of K413R did not regulate the stability of the SF3B and U2AF components. Next, we performed an immunoprecipitation assay to detect the interaction between the U2AF and SF3B complexes using U2AF2-WT- and U2AF2-K413R-expressing cells. Interestingly, U2AF2 K413R interacted strongly with components of the SF3B complex and with U2AF1 compared to that with U2AF2 WT **(Fig. 4K)**. Notably, the interaction of U2AF2 and U2AF1 was not obviously regulated after RNAase treatment, while the interaction of U2AF2 and SF3B complex was reduced following RNase treatment **(Fig. 4K)**. These findings implied the interactions between U2AF1 and U2AF2 were independent of RNA binding, while the interaction of U2AF2 and SF3B complex were dependent on RNA binding. As a component of the U2AF complex, U2AF2 interacts with U2AF1 to form the U2AF complex, which recognizes the 3′ splice site (3’ SS) of U2 introns and recruits U2 snRNP^17^ **(Fig. 4L)**. Taken together, these data suggest that U2AF2 acetylation regulates the formation of the U2AF complex and recruitment of the SF3B complex.

### 3.8. U2AF2 acetylation under cellular stress is not HAT1-dependent

To identify the histone acetyltransferase (HAT) responsible for U2AF2 acetylation under cellular stress, we co-transfected U2AF2 with different HATs, namely CBP, p300, HAT1, P/CAF, KAT5, and KAT8. Of these, HAT1 mainly promoted U2AF2 acetylation **(Fig. 4M)**. Both endogenous and exogenous co-IP results showed an interaction between HAT1 and U2AF2 **(Fig. 4N)**. Further experiments were performed to determine whether U2AF2 acetylation in response to stress is dependent on HAT1. We found that knockdown of *Hat1* reduced U2AF2 acetylation, while *Hat1* OE induced the acetylation **(Fig. 4O)**. Furthermore, U2AF2 acetylation was not affected by enzymatically defective HAT1 **(Fig. 4P)**. Although U2AF2 interacted with HAT1, the interaction was altered weakly under TGF-β1 stimulation **(Fig. 4Q, left panel)**. Consistently, under *Hat1* knockdown conditions as well, TGF-β1 stimulation further reduced U2AF2 acetylation **(Fig. 4Q, right panel)**. Thus, although HAT1 acetylated U2AF2, the decreased acetylation level of U2AF2 by TGF-β1 stimulation was independent of HAT1.

### 3.9. SIRT4 deacetylates U2AF2 at Lys413

As SIRT4 interacted with U2AF2, we intend to figure out the directly role of SIRT4 in U2AF2 acetylation. We found that SIRT4 OE reduced U2AF2 acetylation, whereas SIRT4 knockdown increased its acetylation **(Fig. 5A)**. Additionally, TGF-β1 stimulated the interaction between SIRT4 and U2AF2 in a dose dependent manner **(Fig. 5B)**. Consistently, the TGF-β1-induced decrease in U2AF2 acetylation was blocked by the knockdown of *Sirt4* **(Fig. 5C)**. In contrast, SIRT4 OE further enhanced the TGF-β1-induced U2AF2 deacetylation **(Fig. 5C)**. Compared to U2AF2 WT, U2AF2 K413R blocked the SIRT4 OE-induced deacetylation of U2AF2 as well as the interaction of U2AF2 with U2AF1 and the SF3B complex under TGF-β1 stimulation **(Fig. 5D)**. Accordingly, these phenotypes induced by SIRT4 OE were restricted when an enzyme-defective SIRT4 (SIRT4 H161Y) was used **(Fig. 5E)**. Molecular docking simulations showed that U2AF2 Lys413 was present at the contact surface between SIRT4 and U2AF2 **(Fig. 5F)**, further demonstrating that SIRT4 can interact with U2AF2 to promote its deacetylation.

**Figure 5.**
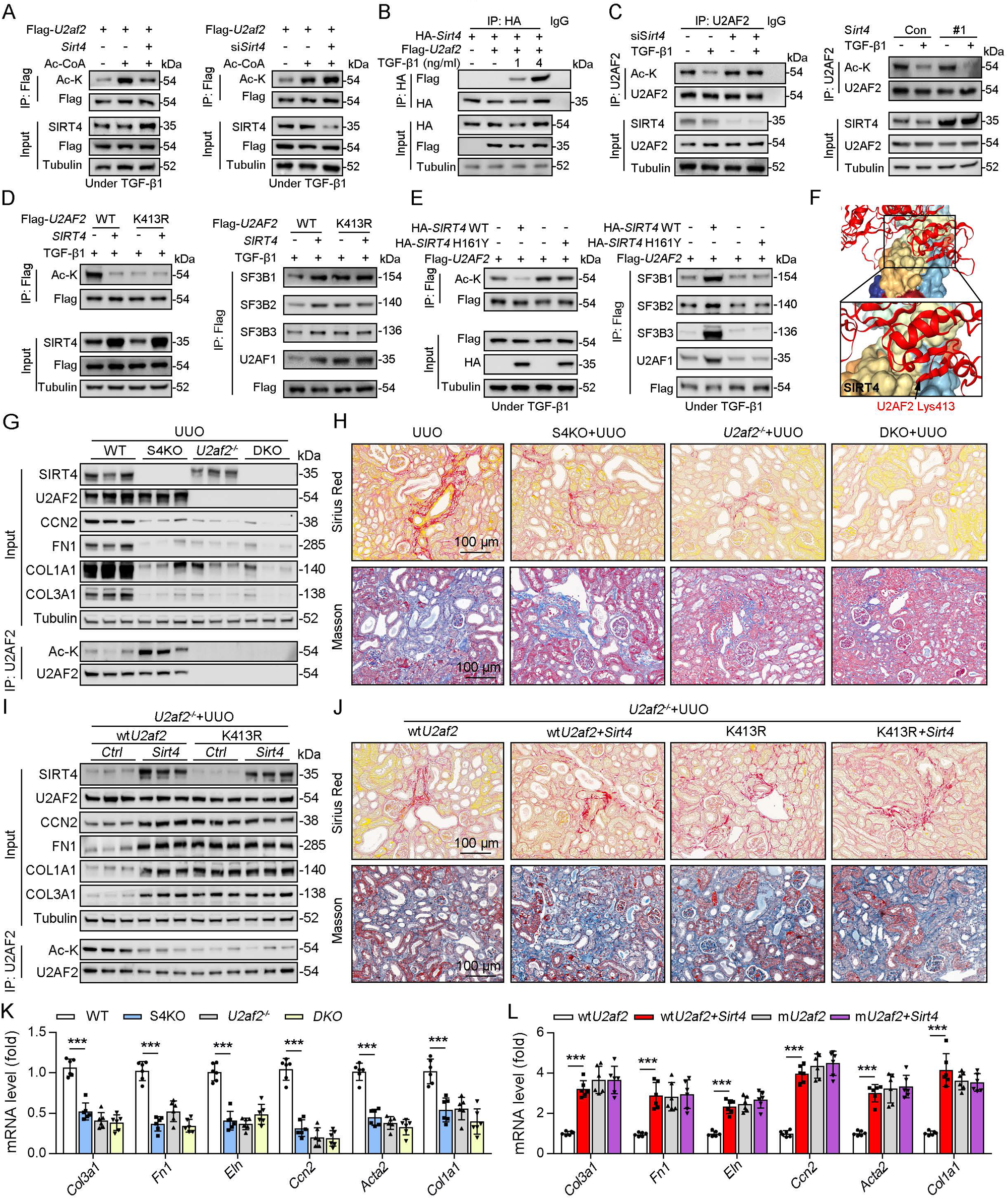
U2AF2 deacetylation under TGF-β1 is SIRT4 dependent. **(A)** WT TECs were transfected with Flag-*U2af2* WT, Ad-*Sirt4* or Ad-sh*Sirt4* under Ac-CoA treatment. Cells lysates subjected to Co-IP with anti-Flag antibody, and Western blotting using indicated antibodies. **(B)** WT TECs were transfected with Flag-*U2af2* WT and HA-*Sirt4* under TGF-β1 stimulation. Cells lysates subjected to Co-IP with anti-HA antibody and Western blotting using indicated antibodies. **(C)** WT TECs were transfected with Ad-sh*Sirt4* or Ad-*Sirt4* with or without TGF-β1 stimulation. Cells lysates subjected to Co-IP with anti-U2AF2 antibody, and Western blotting using indicated antibodies. **(D)** HK2 cells were transfected with Flag-*U2AF2* WT or K413R, Ad-sh*SIRT4* under TGF-β1 stimulation. Cells lysates subjected to Co-IP with anti-Flag antibody, and Western blotting using indicated antibodies. **(E)** HK2 cells were transfected with HA-*SIRT4* WT or H161Y and Flag-*U2AF2* under TGF-β1 treatment. Cells lysates subjected to Co-IP with anti-Flag antibody, and Western blotting using indicated antibodies. **(F)** SIRT4-U2AF2 docking with the HDOCK server. High magnification of boxed areas is presented on the left. Arrow indicates K413 of U2AF2 protein. **(G, H, K)** WT, S4KO, U2AF2^−/−^ or DKO mice were randomly assigned to UUO surgery according to an established protocol. Kidney samples were obtained from mice on day 10 post surgery or sham surgery (n = 6 per group). G: Western blot analysis of the expression of SIRT4, U2AF2, CCN2, FN1, COL1A1, COL3A1, and Tubulin in the kidney from mice. Kidney lysates subjected to Co-IP with anti-U2AF2 antibody, and Western blotting using indicated antibodies. H: Representative images of Masson’s trichrome staining and Sirius red in kidneys from mice (scale bar, 100 μm). K: The mRNA level of *Col1a1, Fn1, Eln, Ccn2, Acta2* and *Col3a1* in the kidney of mice. **(I, J, L)** *U2af2^−/−^* mice received *in situ* renal injection of AAV9-*Ksp-U2af2 WT* (*wtU2af2*), *wtU2af2* plus *Sirt4,* AAV9-*Ksp-U2af2 K413R* (*mutU2af2*), *K413R* plus *Sirt4* at 6 weeks of age. After 2 weeks, the mice were randomly assigned to sham surgery or UUO surgery according to an established protocol. Kidney samples were obtained from mice on day 10 post surgery or sham surgery (n = 6 per group). I: Western blot analysis of the expression of SIRT4, U2AF2, CCN2, FN1, COL1A1, COL3A1, and Tubulin in the kidney from mice. Kidney lysates subjected to Co-IP with anti-U2AF2 antibody, and Western blotting using indicated antibodies. J: Representative images of Masson’s trichrome staining and Sirius red in kidneys from mice (scale bar, 100 μm). L: The mRNA level of *Col1a1, Fn1, Eln, Ccn2, Acta2 and Col3a1* in the kidney of mice. For all panels, data are presented as mean ± SD. \**P* < 0.05, \*\**P* < 0.01, \*\*\**P* < 0.001 by one-way ANOVA with Bonferroni correction test.

To determine whether S4KO alleviates renal fibrosis by upregulating the acetylation of U2AF2 in mouse kidney, we crossed *Cdh16*-cre/ERT2×*U2af2^flox/flox^*(*U2af2*^−/−^) with *Sirt4*^−/−^ mice to generate DKO mice and then performed UUO surgery. No body weight differences were noted due to genetic manipulation (data not shown). Compared to that in WT mice, the acetylation level of U2AF2 was increased, but renal fibrosis was reduced in S4KO mice after UUO surgery **(Fig. 5G, H, K).** The UUO-induced kidney injury was largely reduced in DKO and U2af2^−/−^ mice, which showed comparable collagen deposition and expression of ECM-related proteins and mRNA **(Fig. 5G, H, K)**. We next treated U2af2^−/−^ mice with AAV-wt*U2af2* or AAV-m*U2af2* (*U2af2* K413R) to re-express the two types of U2AF2 proteins through in situ renal injection. Mice were followed by administered of AAV-*Sirt4* or AAV-Ctrl and then subjected to UUO surgery. The U2AF2 K413R OE mice showed a greater extent of renal fibrosis than did the wtU2AF2 OE mice, as evidenced by collagen deposition and increased ECM protein levels **(Fig. 5I, J, L).** Importantly, the extent of renal fibrosis was remarkably augmented in mice injected with wt*U2af2* OE and *Sirt4* OE compared to that in wtU2AF2 OE mice **(Fig. 5I, J, L)**. However, *Sirt4* OE had little synergistic effect on mice injected with AAV-m*U2af2* **(Fig. 5I, J, L)**. Together, these data support that SIRT4-mediated deacetylation of U2AF2 at K413 is an important step in SIRT4-induced renal fibrosis.

### 3.10. SIRT4 deacetylates U2AF2 at K413 to regulate *CCN2* expression and pre-mRNA splicing

U2AF1 S34F and Q157R mutations have been reported to compromise U2AF2–RNA interactions, resulting predominantly in intron retention and exon exclusion^31^. Hence, we hypothesized that the SIRT4-mediated deacetylation of U2AF2 participates in the regulation of these processes. RNA-sequencing was performed to identify the changes in gene expression and alternative splicing in adenovirus-mediated SIRT4^OE^-transfected cells or adenovirus-Ctrl-transfected cells after TGF-β1 stimulation **(Fig. 6A)**. Surprisingly, SIRT4 stimulation upregulated several genes (*P* < 0.01; Table EV1) that are involved in mRNA processing and the TGF-β signaling pathway. A total of 142 differentially expressed genes (*P* < 0.0005) were identified **(Fig. 6A)**. Meanwhile, 248 genes showed differential intron retention (FDR < 0.05; Table EV2) **(Fig. 6A)**. This analysis revealed four genes that were common between the differentially expressed genes and the differentially spliced genes **(Fig. 6A)**. *Ccn2*, one of these four genes, ranked third among the all upregulated genes (*P* < 0.0005) after SIRT4 overexpression. Given that intron retention due to abnormal splicing can trigger nonsense-mediated mRNA decay, gene expression could be affected by a change in splicing pattern. The mRNA levels of *CCN2* were remarkably increased in SIRT4^OE^ cells but reduced in *Sirt4-*knockdown cells **(Fig. 6B)**. In addition, *CCN2* mRNA levels were significantly increased in U2AF2 K413R-transfected cells but remarkably reduced in U2AF2 K413Q [lysine to glutamine (Q) mutant for protein hyperacetylation mimic]-transfected cells **(Fig. 6B)**. This finding prompted us to investigate the role of specific lysine residues within U2AF2 that may be critical for the regulation of CCN2 expression. To this end, we introduced the K453Q mutation as a control to discern the distinct effects of lysine acetylation at different sites. Our results showed that SIRT4 OE significantly elevated the protein levels of CCN2 in U2AF2 WT or U2AF2 K453Q-transfected cells but not in U2AF2-K413Q or K413R-transfected cells **(Fig. 6C, D)**, suggesting that only the U2AF2 acetylation at K413 is efficient to regulate CCN2 expression.

**Figure 6.**
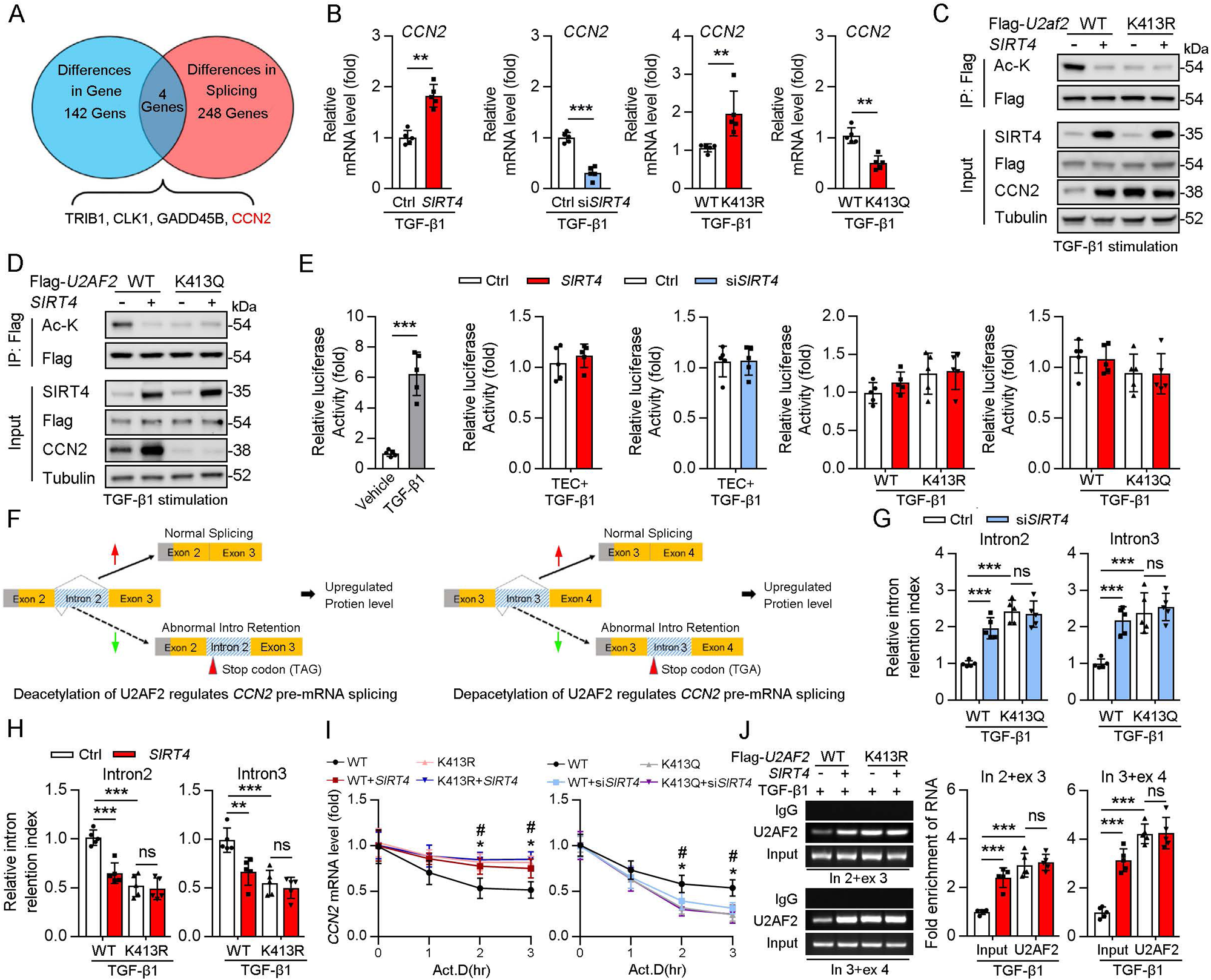
Acetylation of U2AF2 at K413 regulates CCN2 gene expression and pre-mRNA splicing. **(A)** The crosstalk of differences of the gene expression profiling analysis and splicing analysis. **(B)** The *Ccn2* mRNA level in HK2 cells (n = 5). **(C, D)** HK2 cells were transfected with Flag-*U2AF2* WT, K413R, or K413Q, and Ad-*SIRT4* or *SIRT4* siRNA (si*SIRT4*) as indicated in figure. Cells lysates subjected to Co-IP with anti-Flag antibody, and Western blotting using indicated antibodies. **(E)** Luciferase activity of *Ccn2* promoter in HK2 cells transfected with Ad-*SIRT4*, Ad-sh*SIRT4*, WT *U2AF2, U2AF2* K413R, or *U2AF2* K413Q under TGF-β1 stimulation was measured (n = 5). **(F)** Schematics of *Ccn2* alternative splicing pattern regulated by hyperacetylated U2AF2. Full line and dotted line represent normal splicing and abnormal intron retention, respectively. Grey and yellow boxes represent untranslated region (UTR) and protein coding region (CDS), respectively. The first stop codon in intron 2 or 3 was indicated with red arrow. **(G, H)** Quantitative real-time PCR analysis of normal or abnormal splicing isoforms of *CCN2* (n = 5). **(I)** Actinomycin D (Act D, 2 mg/mL) treatment and quantitative real-time PCR were performed to measure *CCN2* mRNA levels (n = 5). **(J)** HK2 cells were transfected with Flag-*U2AF2* WT, Flag-*U2AF2* WT, or Flag-*U2AF2* K413R with or without Ad-*SIRT4* under TGF-β1 stimulation. Cells lysates subjected to RNA Binding Protein Immunoprecipitation (RIP) with anti-U2AF2 antibody (left panel). Quantitative real-time PCR results of RIP assays (right panel). For all panels, data are presented as mean ± SD. \*\**P* <C0.01, \*\*\**P* <C0.001, by Student’s t-test for B. \*\**P* < 0.01, \*\*\**P* < 0.001 by one-way ANOVA with Bonferroni correction test for E, G, H, I and J.

CCN2 is transcriptionally regulated by Sp1 in response to TGF-β1 stimulation^32^. Thus, we examined whether SIRT4 induced CCN2 expression through transcription co-activator function. An analysis of the promoter region of *Ccn2* (including Sp1 binding site) showed that there was no increase in the promoter activity following overexpression of U2AF2 WT, U2AF2-K413R or U2AF2-K413Q upon TGF-β1 stimulation **(Fig. 6E)**, implying that the SIRT4-induced CCN2 expression is not transcription dependent. According to the RNA sequencing data, SIRT4 stimulation increased the efficiency of *Ccn2* pre-mRNA exon2/intron2 and exon3/intron3 splicing **(Fig. 6F)**. Upon analyzing the gene sequence of *Ccn2*, we found that both *Ccn2* introns 2 and 3 contain a TGA stop codon, implying that they could trigger the degradation of abnormal mRNAs. Furthermore, we confirmed that SIRT4 knockdown or U2AF2 K413Q increased the retention index of *Ccn2* introns 2 and 3, whereas SIRT4 OE or U2AF2 K413 reduced the retention index **(Fig. 6G, H)**. U2AF2 K413R or K413Q impeded the changes in intron 2 and 3 retention indices of *Ccn2* caused by SIRT4 OE or *Sirt4* knockdown, respectively **(Fig. 6G, H)**. Next, we assessed whether U2AF2 regulates CCN2 expression by regulating mRNA stability. We treated SIRT4 OE or *Sirt4* knockdown cells with Actinomycin D to block transcription and measured *Ccn2* mRNA stability over time. The mRNA stability of *Ccn2* was significantly enhanced in SIRT4 OE cells, whereas it was reduced in *Sirt4* knockdown cells compared to that in control cells **(Fig. 6I)**. As expected, compared to that in U2AF2 WT-transfected cells, U2AF2 K413R increased the mRNA stability of *Ccn2*, while U2AF2 K413Q reduced its stability **(Fig. 6I)**. In addition, K413R and K413Q repressed the changes in the stability of *Ccn2* mRNA caused by SIRT4 OE or *Sirt4* knockdown, respectively **(Fig. 6I)**. Next, a RiboIP experiment was performed wherein U2AF2 was immunoprecipitated to determine whether it can bind endogenous *Ccn2* transcripts **(Fig. 6J)**. SIRT4 OE enhanced the interaction between U2AF2 and *Ccn2* pre-mRNA (indicated by the PCR product containing intron 2 and exon 3 or intron 3 and exon 4) in U2AF2 WT-transfected cells but had little effect on their interaction in U2AF2 K413R-transfected cells **(Fig. 6J)**. This indicates that the SIRT4-induced deacetylation of U2AF2 at K413 increases CCN2 expression by regulating the alternative splicing of *Ccn2*.

### 3.11. SIRT4 translocates from mitochondria to the cytoplasm through the BAX/BAK pore after TGF-β stimulation

SIRT4 is located in the mitochondrial matrix under normal conditions^33^. A previous study showed that SIRT4 translocates from the mitochondria into the cytoplasm upon Wnt stimulation^34^. Hence, we conducted an immunofluorescence assay and organelle separation experiment to compare the localization of SIRT4 before and after TGF-β1 treatment. Interestingly, SIRT4 localization changed from mitochondria to the cytoplasm or even nucleus at after 12 h of TGF-β1 treatment **(Fig. 7A–C)**. Moreover, TGF-β1 stimulation did not induce U2AF2 release from the nucleus into the cytoplasm, even after 24 h (data not shown). Other stimulations such as serum starvation, tunicamycin, poly (I: C), and cisplatin also induced this translocation of SIRT4 **(Fig. 7D)**. It is well known that BAX and BAK are two key molecules of the mitochondrial permeability transition pore. They form polymers on the mitochondrial outer membrane and mediate the release of mitochondrial contents such as mtDNA, mitochondrial dsRNA, and cytochrome c^35,36^. Therefore, we tested whether the TGF-β1-induced release of SIRT4 is dependent on the BAX/BAK oligomeric pore. Notably, BAX or BAK deficiency or MSN-125 (an effective oligomeric inhibitor of Bax and Bak) treatment almost abolished the release of SIRT4 from mitochondria to the cytoplasm as well as the upregulation of *Ccn2* under TGF-β1 stimulation **(Fig. 7E, F)**. As expected, the *in vivo* results showed that MSN-125 inhibited the translocation of SIRT4 from mitochondria to the cytoplasm in the kidneys of mice following UUO surgery **(Fig. 7G)**. Together, these results suggest that TGF-β1 induces the translocation of SIRT4 from mitochondria to the cytoplasm in a BAX/BAK-dependent manner.

**Figure 7.**
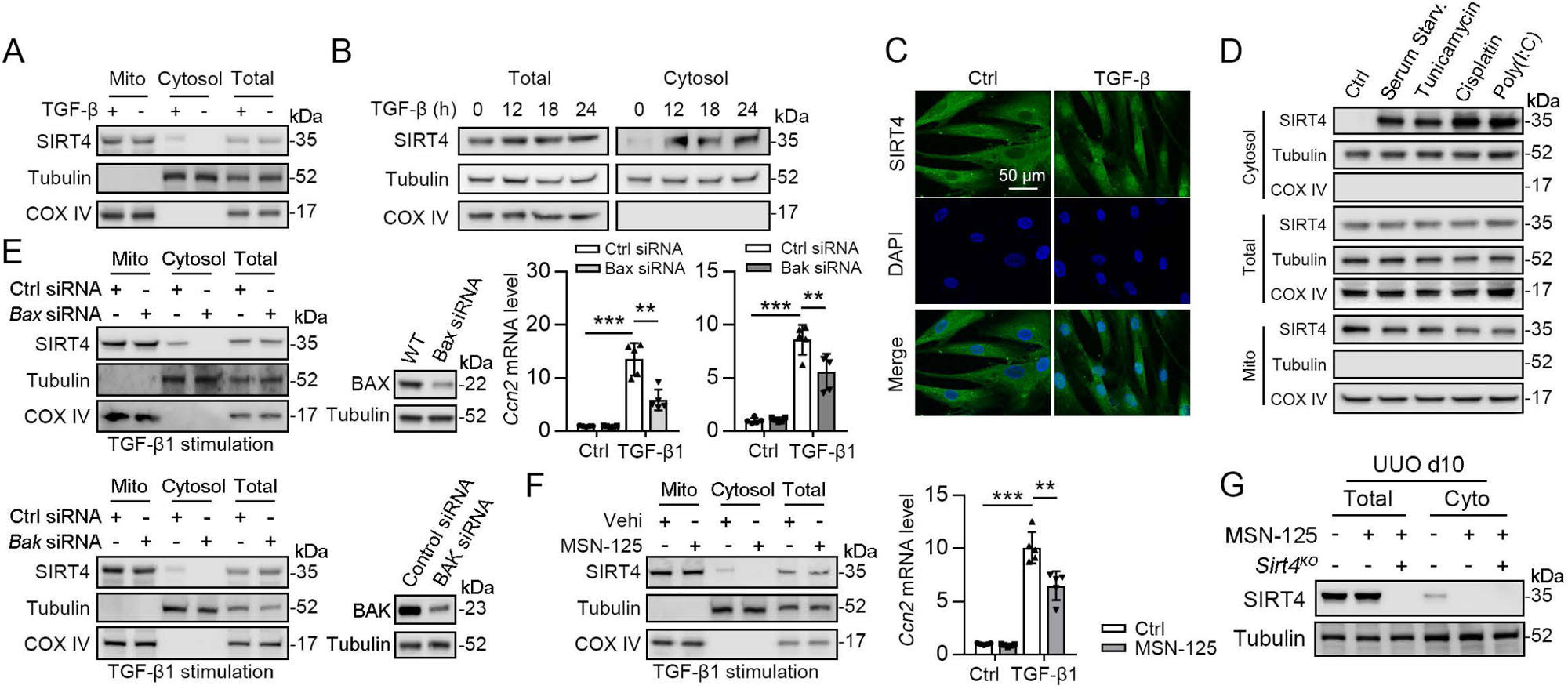
SIRT4 is released through BAX/BAK pore in a TGF-β1-dependent manner. **(A)** Organelle separation experiment and immunoblot analysis detected the expression/localization of SIRT4, Tubulin, and COX IV. **(B)** TECs were treated with TGF-β1 for indicated time. Organelle separation experiment and immunoblot analysis detected the localization of SIRT4, Tubulin, and COX IV. **(C)** Representative image of immunofluorescent staining of SIRT4 in TECs treated with TGF-β1 or not (scale bar, 50 μm). **(D)** TECs were treated with Tunicamycin, Cisplatin, Poly(I:C), or under serum starvation. Organelle separation experiment and immunoblot analysis detected the localization of SIRT4, Tubulin, and COX IV. **(E)** WT TECs incubated with *Bax* siRNA or *Bak* siRNA under TGF-β1 stimulation. Organelle separation experiment and immunoblot analysis detected the localization of SIRT4, Tubulin, and COX IV (left panels). Western blot analysis of the expression of BAX, BAK and Tubulin in TECs (middle panels). The mRNA level of *Ccn2* was detected by qPCR (n = 5) (right panels). **(F)** WT TECs were treated with MSN-125 (10 μM) or vehicle. Organelle separation experiment and immunoblot analysis detected the localization of SIRT4, Tubulin, and COX IV (left panel). The mRNA level of *Ccn2* was detected by qPCR (n = 5) (right panel). **(G)** Kidney tissues from WT or S4KO mice were treated with MSN-125 or vehicle. The mRNA level of *Ccn2* was detected by qPCR in the kidney of mice (n = 5). For all panels, data are presented as mean ± SD. \*\**P* < 0.01, \*\*\**P* < 0.001 by one-way ANOVA with Bonferroni correction test.

### 3.12. ERK phosphorylates SIRT4 at Ser36 to promote the binding of SIRT4 to importin α1 and nuclear translocation

As SIRT4 showed nuclear localization **(Fig. 7C)** and interacted with the nuclear protein U2AF2 **(Fig. 4C)** under TGF-β1 stimulation, we investigated the mechanisms underlying SIRT4 accumulation in the nucleus. Pretreatment of TECs with LY290042 (phosphoinositide 3-kinase inhibitor), SU6656 (Src inhibitor), SP600125 (JNK inhibitor), and U0126 (MEK/ERK inhibitor) blocked TGF-β1-induced phosphorylation of AKT, c-Src, c-Jun, and ERK1/2, respectively. Immunoblotting analyses showed that only U0126 treatment abrogated the TGF-β1-induced nuclear translocation of SIRT4 **(Fig. 8A, upper panel)**. Compared to the amount of cytosolic SIRT4, nuclear SIRT4 is present in a small portion **(Fig. 8A, bottom panel)**. These results were further supported by the immunofluorescence analyses **(Fig. 8B)**. Additionally, expression of the Flag-ERK2 K52R kinase-dead mutant blocked the TGF-β1-induced nuclear accumulation of SIRT4 and resulted in the accumulation of phosphorylated SIRT4 in the cytosol **(Fig. 8C, left panel)**. Co-expression of a constitutively active MEK1 Q56P mutant (expression of constitutively active MEK1 Q56P with WT ERK2) with Flag-tagged ERK2 WT or ERK2 K52R in TECs **(Fig. 8C, right panel)** showed that expression of WT ERK2, but not ERK2 K52R, induced the nuclear translocation of SIRT4. These results indicate that ERK activation is required for the TGF-β1-induced nuclear translocation of SIRT4.

**Figure 8.**
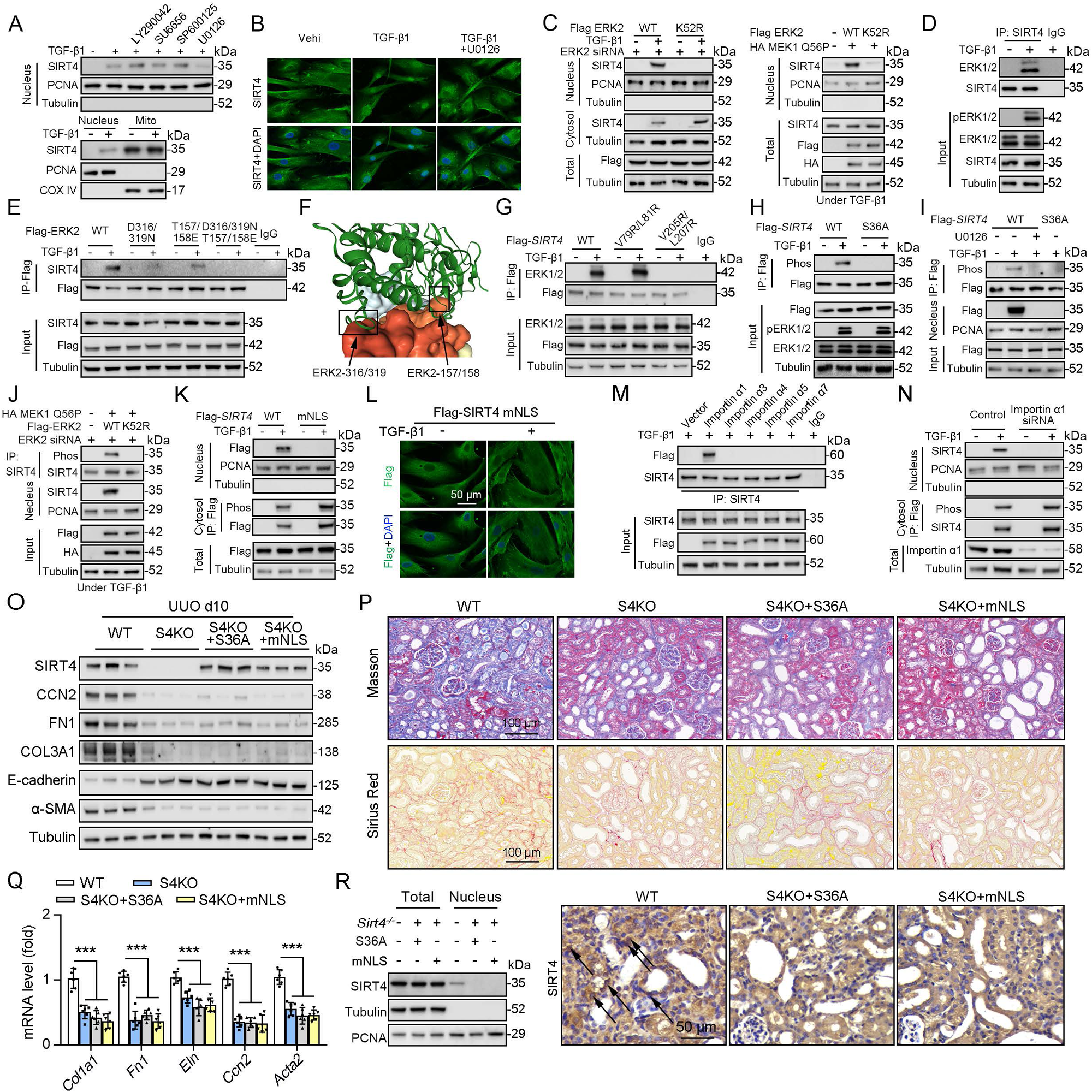
ERK2 phosphorylates SIRT4 at S36 and promotes it nucleus translocation. **(A)** (The upper panel) Nuclear fractions were prepared from TECs pretreated with LY290042 (30 μM), SU6656 (4 μM), SP600125 (25 μM), and U0126 (20 μM) for 30 min before TGF-β1 (2 ng/ml) for 12 h. Nuclear PCNA and cytoplasmic Tubulin were used as controls. (The bottom panel) TECs were treated with TGF-β1 (2 ng/ml) for 12 h. Organelle separation experiment and immunoblot analysis detected the localization of SIRT4, PCNA, and Tubulin. **(B)** TECs were pretreated with or without U0126 (20 μM) for 30 min, then treated with TGF-β1 (2 ng/ml) for 12 h. Immunofluorescence analyses were performed with the indicated antibodies. **(C)** HK2 cells were stably transfected with Flag-ERK2, Flag-ERK2 K52R, or siRNA ERK2 (left panel) or transiently transfected with HA-MEK1 Q56P and indicated Flag-tagged ERK2 proteins (right panel). The cells were treated with or without TGF-β1 (2 ng/ml) for 12 h, and the total cell lysates, cytosol and nuclear fractions were prepared for determination of indicated proteins by Western blot. **(D)** TECs were treated with TGF-β1, and cells lysates subjected to Co-IP with anti-SIRT4 antibody, and Western blotting using indicated antibodies. **(E)** HK2 cells transfected with vectors expressing the indicated Flag-tagged ERK proteins were treated with or without TGF-β1 (2 ng/ml) for 12 h. **(F)** SIRT4-ERK2 docking with the HDOCK server. **(G)** HK2 cells expressing the indicated Flag-*SIRT4* proteins were treated with or without TGF-β1 (2 ng/ml) for 12 h. **(H, I)** HK2 cells transfected with Flag-Sirt4 WT or S36A and then incubated with TGF-β1 and/or U0126 as indicated in figure, the total cell lysates and nuclear fractions were prepared. Total cells lysates subjected to Co-IP with anti-Flag antibody, and Western blotting using indicated antibodies. **(J)** HK2 cells were transfected with MEK1 Q56P, Flag-ERK2 WT or K52R, and ERK2 siRNA as indicated in figure, and the total cell lysates and nuclear fractions were prepared. Total cells lysates subjected to Co-IP with SIRT4 antibody and Western blotting using indicated antibodies. **(K, L)** HK2 cells transfected with Flag-*Sirt4* WT or mNLS with or without TGF-β1 treatment, and the total cell lysates, cytosol, and nuclear fractions were prepared. Western blotting using indicated antibodies (K). Immunofluorescence analyses were performed with the indicated antibodies (L). **(M)** The indicated Flag-importin α proteins were transfected in HK2 cells, then treated with TGF-β1 (2 ng/ml) for 12 h. Flag-importin α proteins were immunoprecipitated with an anti-Flag antibody. **(N)** HK2 cells transfected with importin α1 were treated with or without TGF-β1 (2 ng/ml) for 12 h. The total cell lysates, cytosolic and nuclear fractions were prepared and Western blotting using indicated antibodies. **(O to R)** S4KO mice received *in situ* renal injection of AAV9-*Ksp-Sirt4 S36A* (S36A) or AAV9-*Ksp-Sirt4 mNLS* (mNLS) at 6 weeks of age. After 2 weeks, the mice were randomly assigned to sham surgery or UUO surgery according to an established protocol. O: Western blot analysis of the expression of SIRT4, CCN2, FN1, COL3A1, E-cadherin, α-SMA and Tubulin in mouse kidney. P: Representative images of Masson’s trichrome staining and Sirius red staining in kidney sections from mice (scale bar, 100 μm). Q: The mRNA level of *Col1a1, Fn1, Eln, Ccn2 and Acta2* in the kidney of mice. R: The total cell lysates and nuclear fractions were prepared from kidney and Western blotting using indicated antibodies (left panel). (Right panel) Representative images of immunohistochemical staining of SIRT4 in kidneys from mice (scale bar, 100 μm). For all panels, data are presented as mean ± SD. \*\*\**P* < 0.001 by one-way ANOVA with Bonferroni correction test.

To further determine the relationship between ERK1/2 and SIRT4, we performed a Co-IP assay and found that TGF-β1 treatment resulted in the binding of ERK1/2 to SIRT4 **(Fig. 8D)**. MAP kinases bind to their substrates through a docking groove comprising an acidic common docking (CD) domain and glutamic acid-aspartic acid (ED) pockets^37^. Our results showed that mutation of either the ERK2 CD domain (D316/319N) or the ED pocket (T157/158E) reduced the binding to endogenous SIRT4 compared to that in the WT ERK2 control **(Fig. 8E)**. Combined mutations at both the CD domain and ED pocket (T/E-D/N) abrogated the binding of ERK2 to SIRT4 entirely **(Fig. 8E)**, indicating that ERK2 binds SIRT4 through its docking groove. Furthermore, the server provided docking information regarding SIRT4–ERK2, indicating SIRT4 and ERK2 CD domain and ED pocket interactions **(Fig. 8F)**. ERK substrates often have a docking domain characterized by a cluster of basic residues, followed by an LXL motif (L represents Leu, but can also be Ile or Val; X represents any amino acid)^37^. An analysis of the docking information of SIRT4 - ERK2 and the amino acid sequence of SIRT4 revealed the putative ERK-binding sequence 77-EKVGLYARTDRR-88 and 203-GDVFLSE-209, which contain LXL motifs at V79/L81 and V205/L207, respectively. Immunoblotting of the immunoprecipitated Flag-SIRT4 proteins with an anti-ERK1/2 antibody showed that a SIRT4 V205/L207 mutant, but not a SIRT4 V79/L81 mutant, markedly reduced its binding to ERK1/2 **(Fig. 8G)**. These results indicate that the ERK2 docking groove binds to a CD domain in SIRT4 at V205/L207. Sequence analysis of SIRT4 revealed that it contains an ERK consensus phosphorylation motif (Ser-Pro) at the S36/P37 residues. Notably, S36A mutation completely abrogated the TGF-β1-dependent phosphorylation of SIRT4 **(Fig. 8H)**. Consistently, pretreatment with U0126 blocked the TGF-β1-induced S36 phosphorylation and nuclear translocation of SIRT4 **(Fig. 8I)**. In addition, expression of constitutively active MEK1 Q56P with WT ERK2, but not of ERK2 K52R, induced SIRT4 phosphorylation **(Fig. 8J)**. These results indicate that ERK2 specifically phosphorylates SIRT4.

Notably, we found that SIRT4 contains a potential nuclear localization signal (NLSs), a single type containing 3–5 basic amino acids with the weak consensus Lys-Arg/Lys-X-Arg/Lys^38^, 248-KRVK-251. We mutated the K248/251 and R249 residues in the putative NLS sequences of SIRT4 to alanine (named as mNLS). Cell fractionation and immunofluorescence analyses showed that Flag-SIRT4-mNLS, unlike the WT SIRT4, was unable to translocate into the nucleus upon TGF-β1 treatment **(Fig. 8K, L)**. These results indicate that the NLS in SIRT4 is essential for TGF-β1-induced nuclear translocation of SIRT4. Importin α functions as an adaptor and links NLS-containing proteins to importin β, which then docks the ternary complex at the nuclear-pore complex to facilitate the translocation of these proteins across the nuclear envelope^39,40^. Six importin α family members (α1, α3, α4, α5, α6 and α7) have been identified in humans^40^. We found that the endogenous SIRT4 only binds importin α1 **(Fig. 8M)**. Importin a1/Rch1 is barely detectable in the glomeruli of normal SD rat kidneys but is highly expressed in tubular cells, and the importin α1/Rch1 staining is significantly enhanced in the kidneys of diabetic rats^41^. Depletion of importin α1 with *Rch1* (coding for importin α1) shRNA **(Fig. 8N)** largely blocked the TGF-β1-induced nuclear translocation of SIRT4 and resulted in the accumulation of phosphorylated SIRT4 in the cytosol **(Fig. 8N)**. *In vivo*, U0126 restrained the *Sirt4^OE^*-induced nuclear translocation of SIRT4 **(Fig. S3A, B)**. Furthermore, SIRT4 S36A or SIRT4 mNLS overexpression failed to accumulate SIRT4 in the nucleus and aggravated renal fibrosis in S4KO UUO mice **(Fig. 8O–R)**. Taken together, these results strongly suggest that TGF-β1-induced SIRT4 nuclear translocation is mediated by SIRT4 phosphorylation at S36, which is regulated by the ERK1/2 signaling pathway.

### 3.13. Exosomes containing anti-SIRT4 antibodies alleviate renal fibrosis in UUO mice

SIRT4 is necessary for maintaining mitochondrial function^8,42^. Since knockout or complete inhibition of SIRT4 may not be an advisable therapeutic strategy, we constructed exosomes containing anti-SIRT4 antibodies (αSRIT4) to treat UUO mice. Exosomes containing αSRIT4 effectively inhibited renal fibrosis in UUO mice, accompanied by a decrease in SIRT4 expression in the nucleus, with little effect on mitochondria SIRT4 content **(Fig. S4A-D)**.

Reportedly, the initial stage of the canonical Wnt signaling pathway in which SIRT4 translocates from mitochondria into the cytoplasm leads to β-catenin protein accumulation^34^. To determine whether the β-catenin accumulation is involved in the SIRT4-mediated kidney fibrosis *in vivo*, we generated mice expressing SIRT4^OE^ TECs and subjected them to UUO surgery with or without MSAB treatment (MSAB binds to β-catenin and promotes its degradation). Our results showed that SIRT4 OE remarkably aggravated renal fibrosis under MSAB treatment **(Fig. S5A–C)**. Moreover, downregulation of β-catenin accumulation by MSAB can inhibit renal fibrosis in WT mice, but not the SIRT4^OE^ mice **(Fig. S5A–C).** These findings suggest that SIRT4-mediated the pathogenesis of renal fibrosis is independent of β-catenin accumulation.

## 4. Discussion

Renal fibrosis, especially tubulointerstitial fibrosis, is an inevitable common pathway of progressive chronic kidney disease^43,44^. However, there is a lack of information regarding the pathogenesis of renal fibrosis, which hampers the development of effective therapeutics^45^. Here, we demonstrate that the nuclear translocation of SIRT4 is a prime initiator of kidney fibrosis. SIRT4 significantly accumulates in the nucleus during fibrosis following obstructed nephropathy and renal ischemia reperfusion injury. Global knockout or target deletion of *Sirt4* in TECs attenuated UUO-induced kidney fibrosis, whereas TEC-specific SIRT4^OE^ aggravated the fibrosis. Mechanistically, we found that TGF-β1 promoted SIRT4 release from mitochondria through the BAX/BAK pore. Furthermore, TGF-β1 activation resulted in the nuclear translocation of SIRT4, which was mediated by the ERK1/2-dependent phosphorylation of SIRT4 at S36, and consequently the binding of SIRT4 to importin α1. Nuclear SIRT4 deacetylates U2AF2 and promotes U2 snRNP formation, which promotes the *Ccn2* pre-mRNA splicing, ultimately leading to the increased CCN2 expression. In vivo, SIRT4 S36 or NLS mutants blocked SIRT4^OE^-aggravated kidney fibrosis in UUO mice, which implied that SIRT4 nuclear translocation plays a significant role in the progression of kidney fibrosis.

Acetyl-CoA is an important energy-rich metabolite for homeostasis. In normal states, abundant acetyl-CoA in the cytosol shuttles freely in the nucleus or mitochondria to modulate the acetylation of histone or non-histone proteins. A rapid reduction in total acetyl-CoA levels in renal cells is observed after TGF-β1 stimulation^46^. Non-enzymatic acetylation levels are strongly reduced by acetyl-CoA. Under such circumstances, few exceptional hyperacetylated proteins can function as key regulators of kidney fibrosis development. In accordance with this hypothesis, we found that U2AF2 acetylation decreased and U2 snRNP formation increased after deacetylation of U2AF2 **(Fig. 4H–K)**. Strikingly, U2AF2-K413, the deacetylation form, promotes the pre-mRNA splicing and expression of *Ccn2* **(Fig. 6C, D)**, supporting the idea that acetylation of U2AF2 is responsible for fibrotic reaction under TGF-β1. Additionally, a recent study showed that U2AF2 can directly bind and stabilize circNCAPG, which participates in the nuclear translocation of ras responsive element binding protein 1, thereby activating the TGF-β pathway and promoting glioma progression^47^. Taking this into consideration, we speculate that U2AF2 may be a positive feedback regulator of TGF-β1, although further investigations are required to ascertain this.

SIRT4 is mainly located in the mitochondria and participates in various mitochondrial metabolic processes^13^. Some studies have revealed a potential role of SIRT4 in fibrosis^48,49^. In heart, loss of SIRT4 has been found to result in the development of fibrosis. These studies indicated the protective role of SIRT4 in mitochondria. In recent research, the authors found that SIRT4 abolishment can ameliorate CCl4-induced hepatic encephalopathy phenotypes, which was mediated by downregulating and detoxifing ammonia through the urea cycle^50^. In our study, we indicated that SIRT4 translocates from the mitochondria to the cytoplasm, a process caused by TGF-β induced mitochondrial damage **(Fig. 7)**. Furthermore, the cytoplasmic SIRT4 was phosphorylated by ERK2, which is a downstream of TGF-β signaling **(Fig. 8A-J)**. The phosphorylated SIRT4 further translocated to the nuclear for promoting CCN2 expression by regulating the alternative splicing, then accelerated the kidney fibrosis **(Fig. 5)**. Our study introduces a novel concept in the field, demonstrating the nuclear translocation of SIRT4 is a key initiator of kidney fibrosis. This finding diverges from previous studies that have primarily focused on SIRT4’s mitochondrial roles, highlighting a new dimension of SIRT4’s function in renal pathophysiology.

Some studies have shown that SIRT4 exerts a protective effect on podocytes. SIRT4 OE prevents glucose-induced podocyte apoptosis and ROS production, thereby alleviating diabetic kidney disease (DKD)^51^. Furthermore, FOXM1 transcriptionally activates SIRT4 and inhibits NF-κB signaling and the expression of the NLRP3 inflammasome to alleviate kidney injury and podocyte pyroptosis in DKD^52^. In the present study, we suggest a profibrotic role of SIRT4 in TECs, which was contributed by upregulated expression of CCN2. Although these are complicatedly related to one another in CKD pathophysiology, injury to TECs is considered a core element that initiates progressive fibrosis^53^. Therefore, we suggested that SRIT4 may perform different roles in different types of cells or subcellular organelles. Moreover, further studies on the role of SIRT4 in DKD are needed to evaluate the safety of anti-SIRT4 therapy.

Overall, our study reveals that TGF-β1 activation resulted in the nuclear translocation of SIRT4, mediated by the ERK1/2-dependent phosphorylation of SIRT4 at S36, and consequently the binding of SIRT4 to importin α1. In the nucleus, SIRT4-mediated U2AF2 deacetylation at K413, a key protein for the spliceosome, acts as a responder under TGF-β1 stimulation. SIRT4 promotes CCN2 expression through alternative pre-mRNA splicing by deacetylating U2AF2, which contributes to the progression of kidney fibrosis. These findings expand the field of epigenetic regulation of fibrogenic gene expression and provide a potential therapeutic target for kidney fibrosis.

## Supporting information

Revised Manuscript

## Non-standard abbreviations

CCN2: Cellular Communication Network Factor 2
TECs: Tubular epithelial cells
Sirts: Sirtuins
DKD: Diabetes kidney disease
U2AF1/U2AF2: U2 Small Nuclear RNA Auxiliary Factor 1 or 2
mito-STAR: Mitochondrial sirtuin 4 tripartite abundance reporters
S4KO: *Sirt4* knockout
UUO: Unilateral ureteral occlusion
uIRI: Unilateral renal ischemia-reperfusion injury
FA: Folic acid
Ngal: Neutrophil gelatinase-associated lipocalin
Kim-1: Kidney injury molecule 1
MPC: Mouse podocytes
GEC: Mouse glomerular endothelial cells
MF: Mouse fibroblast
TEC: Tubule epithelial cells
OE: Overexpression
RIME: Rapid immunoprecipitation and mass spectrometry of endogenous proteins
WT: Wild type
ER: endoplasmic reticulum
snRNPs: Small nuclear ribonucleoproteins
HAT: Histone Acetyltransferase
CD: Common docking

## Acknowledgments

This work was supported by grants from the National Natural Science Foundation of China (82370876 to **Shu Yang**, 82170842 and 82371572 to **Zhen Liang**, 82171556 to **Lin Kang**), Shenzhen Sustainable Development Science and Technology Special Project, China (No. KCXFZ20201221173600001 to **Zhen Liang**), Key Program Topics of Shenzhen Basic Research, China (No. JCYJ20220818102605013 to **Lin Kang**). Sequencing service was provided by Bioyi Biotechnology Co., Ltd. Wuhan, China.

## Contribution statement

**Guangyan Yang**, **Jiaqing Xiang, Lixing Li,** and **Yanchun Li** conducted experiments. **Xiaoxiao Yang** contributed to the acquisition of data, analysis and interpretation of data. **Shu Yang** and **Zhen Liang** drafted the work or revised it critically for important intellectual content. **Lin Kang and Xiaoxiao Yang** analysed the data and revised the article critically for important intellectual content. **Shu Yang** contributed to the conception and the study design. All authors gave their approval of the version to be published.

---

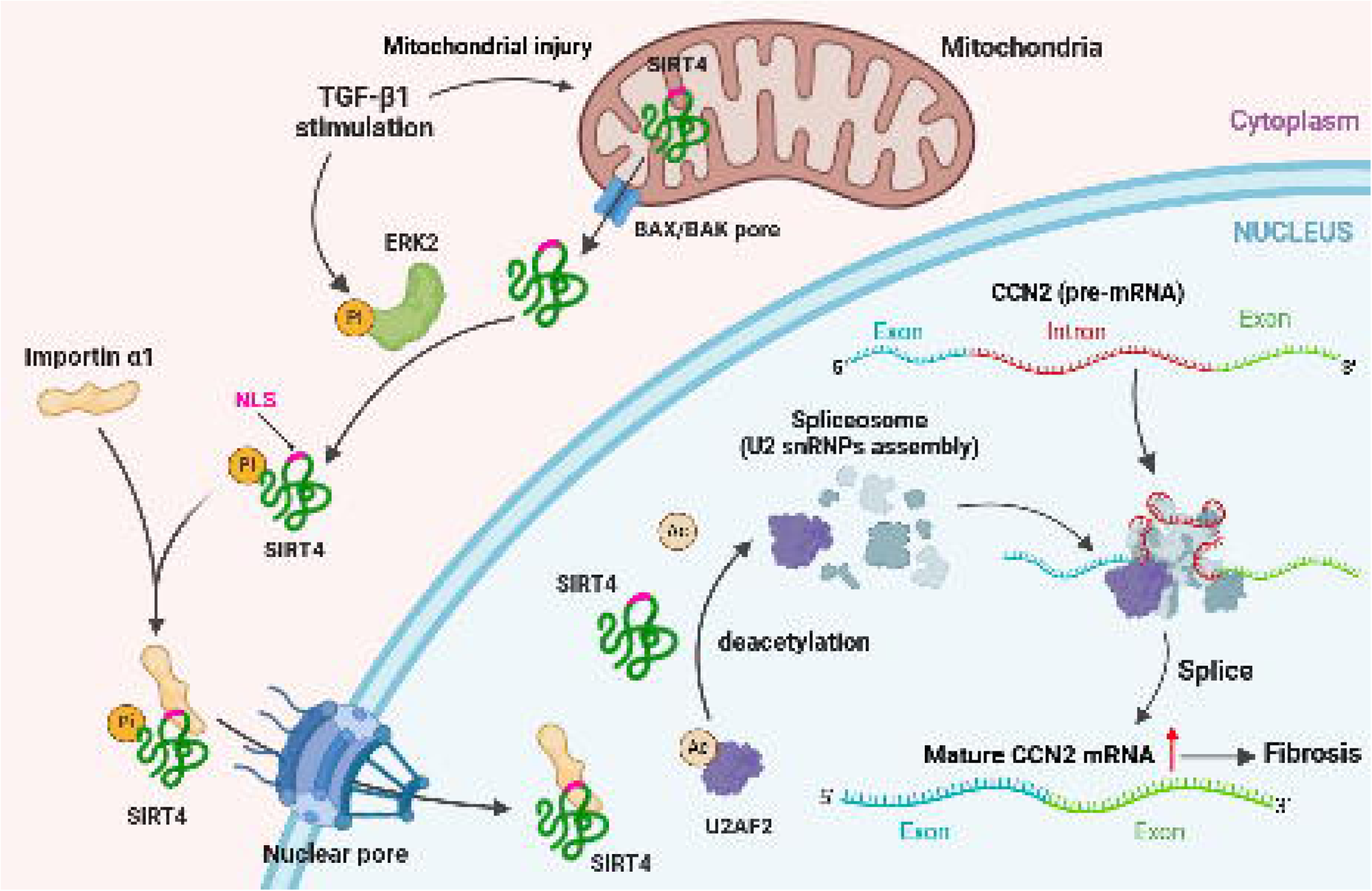

