## Supplementary material for "Nuclear Translocation of SIRT4 Mediates Deacetylation of U2AF2 to Modulate Renal Fibrosis Through Alternative Splicing-mediated Upregulation of CCN2": Revised Manuscript: Electronic supplementary material.docx

**1. ESM Figure and Figure legend**

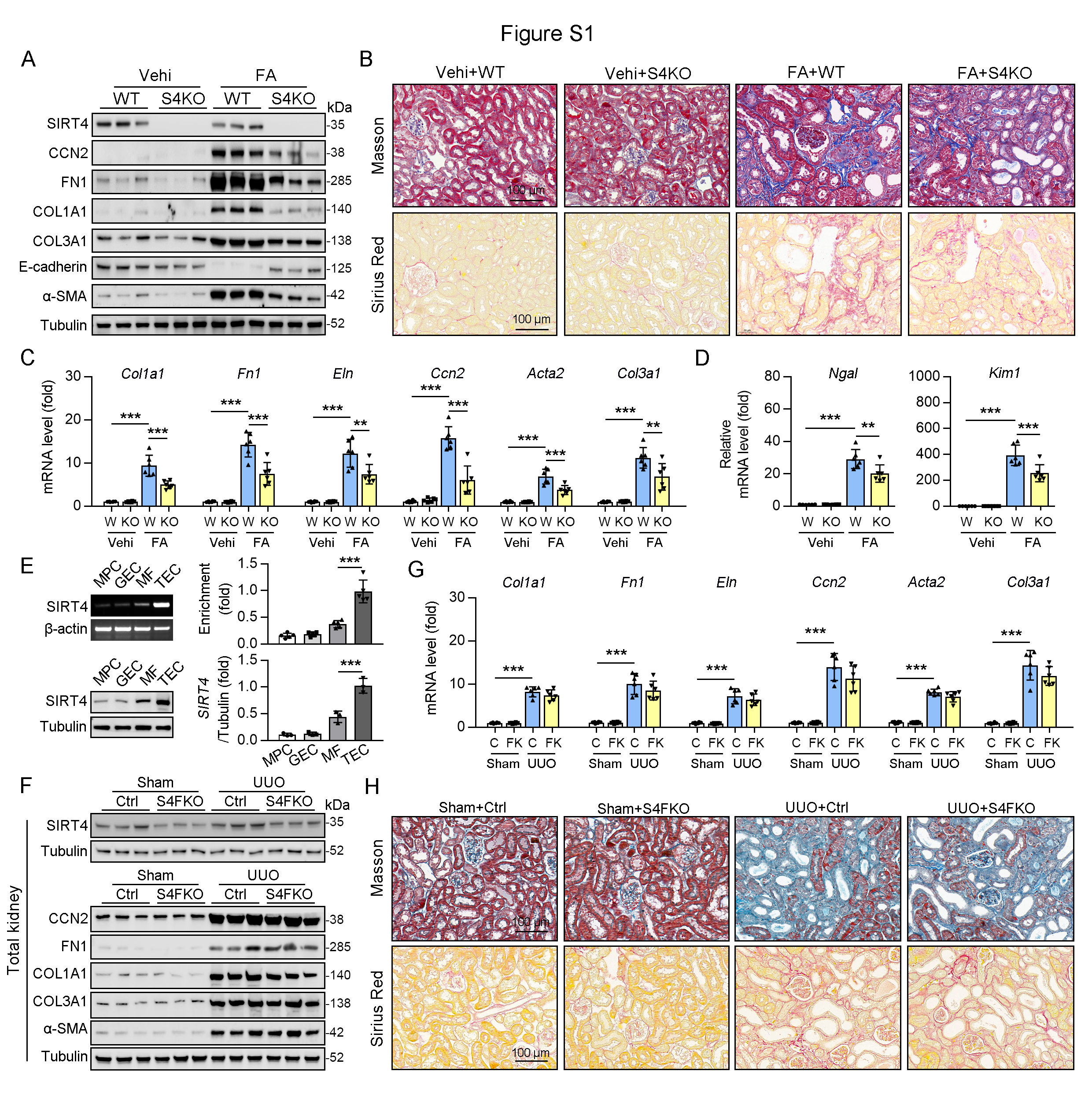

**Figure S1. Knockout of SIRT4 aggravated FA-induced kidney fibrosis. (A to D)** WT or S4KO mice were randomly assigned to FA or vehicle according to an established protocol. Kidney samples were obtained from mice on day 14 post FA or vehicle treatment (n = 6 per group). A: Western blot analysis of the expression of SIRT4, CCN2, FN1, COL1A1, COL3A1, E-cadherin, α-SMA, and Tubulin in the kidney of mice. B: Representative images of Masson’s trichrome staining and Sirius red in kidneys from mice (scale bar, 100 μm). C, D: The mRNA level of *Col1a1, Fn1, Eln, Ccn2, Acta2*, *Col3a1, Ngal*, and *Kim-1* in the kidney of mice*.* **(E)** RT-PCR (n = 5) and western blot analysis (n = 3) of the expression of SIRT4 in selected mouse renal cells including mouse podocytes (MPC), glomerular endothelial cells (GEC), mouse TECs and mouse fibroblast (MF). **(F to H)** Control or S4FKO mice were randomly assigned to sham or UUO surgery according to an established protocol. F: Western blot analysis of the expression of SIRT4, CCN2, FN1, COL1A1, COL3A1, α-SMA, and Tubulin in the kidney of mice. G: The mRNA level of *Col1a1, Fn1, Eln, Ccn2, Acta2*, *Col3a1, Ngal*, and *Kim-1* in the kidney of mice. H: Representative images of Masson’s trichrome staining and Sirius red in kidney sections of mice (scale bar, 100 μm). For all panels, data are presented as mean ± SD. **P* < 0.05, ***P* < 0.01, ****P* < 0.001 by one-way ANOVA with Bonferroni correction test. For all panels, data are presented as mean ± SD. ***P* < 0.01, ****P* < 0.001 by one-way ANOVA with Bonferroni correction test.

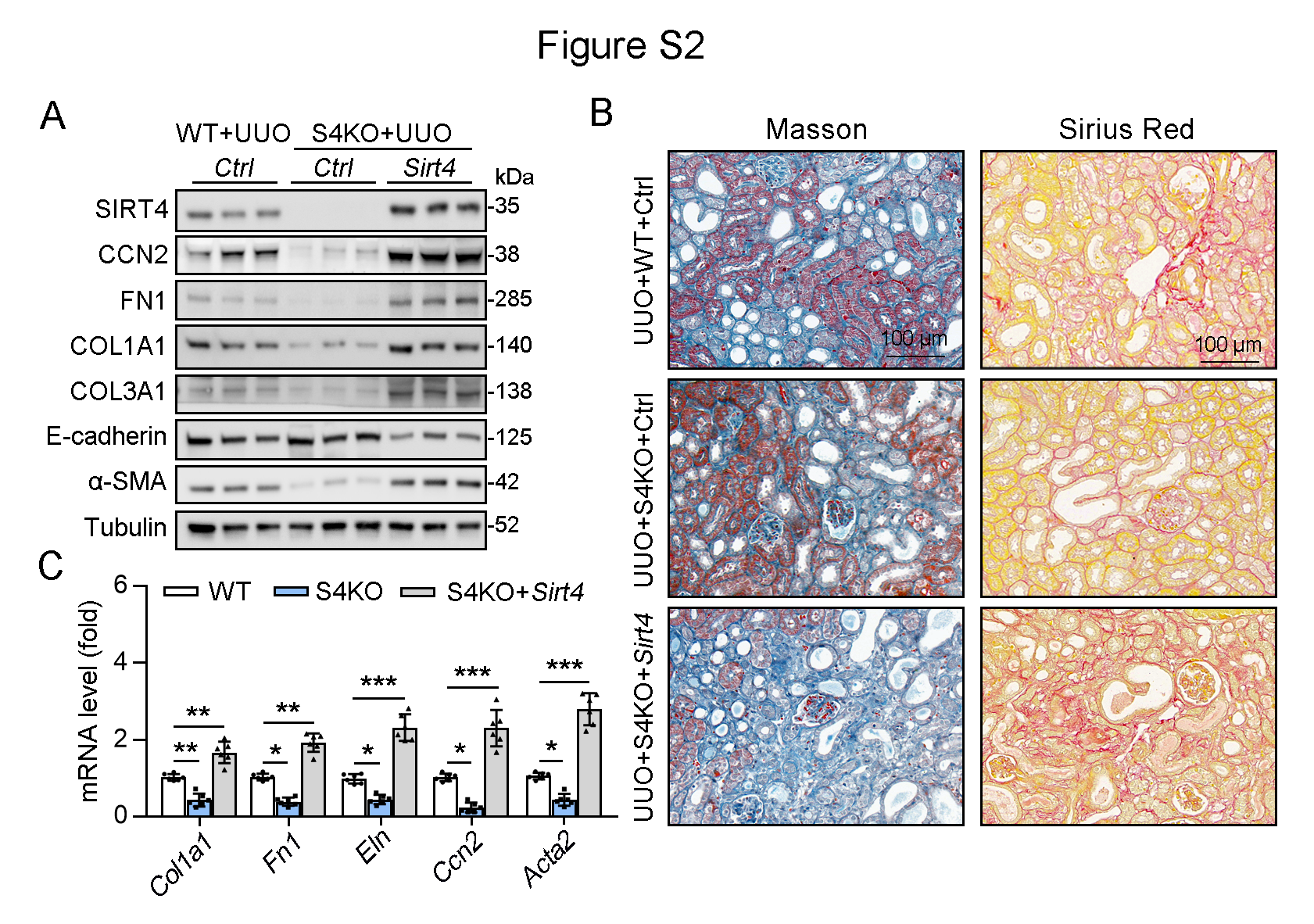

**Figure S2. TECs targeted overexpression of SIRT4 reverses S4KO reduced kidney fibrosis in UUO mice. (A to C)** AAV9-*Ksp*-*Sirt4* was injected into kidneys of mice *in situ* at three independent points in situ (AAV9-*Ksp*-null used as control). After 2-week transfection, the mice received UUO surgery (WT mice treated with AAV9-Ctrl followed by UUO surgery as control) (n = 6 per group). A: Western blot analysis of the expression of SIRT4, CCN2, FN1, COL1A1, COL3A1, E-cadherin, α-SMA, and Tubulin in the kidney of mice. B: Representative images of Masson’s trichrome staining and Sirius red in kidney sections of mice (scale bar, 100 μm). C: The mRNA level of *Col1a1, Fn1, Eln, Ccn2,* and *Acta2.* For all panels, data are presented as mean ± SD. **P* < 0.05, ***P* < 0.01, ****P* < 0.001 by one-way ANOVA with Bonferroni correction test.

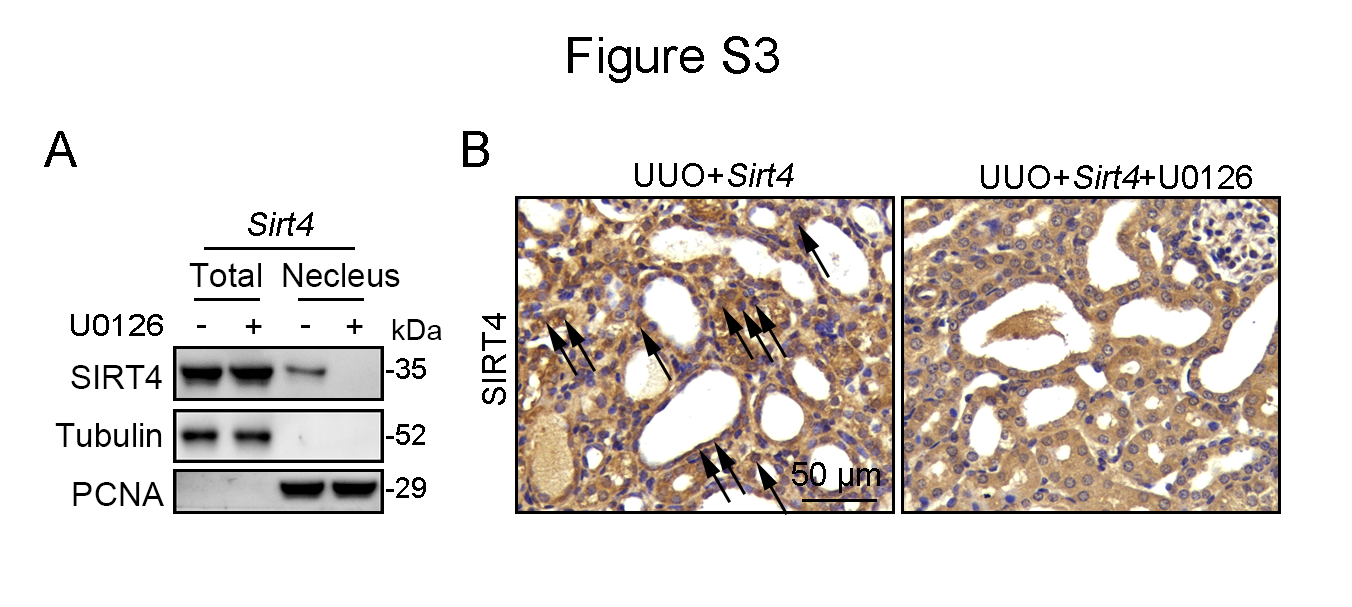

**Figure S3. U0126 prevents SIRT4 overexpression induced kidney fibrosis in UUO mice. (A, B)** AAV9-*Ksp*-*Sirt4* was injected into kidneys of mice *in situ* at three independent points *in situ*. After 2-week transfection, the mice received UUO surgery, and then mice were treated with U0126 (10 mg/kg body weight) or vehicle for 10 days (n = 6 per group). A: Organelle separation experiment and immunoblot analysis detected the localization of SIRT4 and Tubulin in the kidneys of mice. B: Representative images of immunohistochemical staining of SIRT4 (scale bar, 50 μm) in the kidney sections of mice.

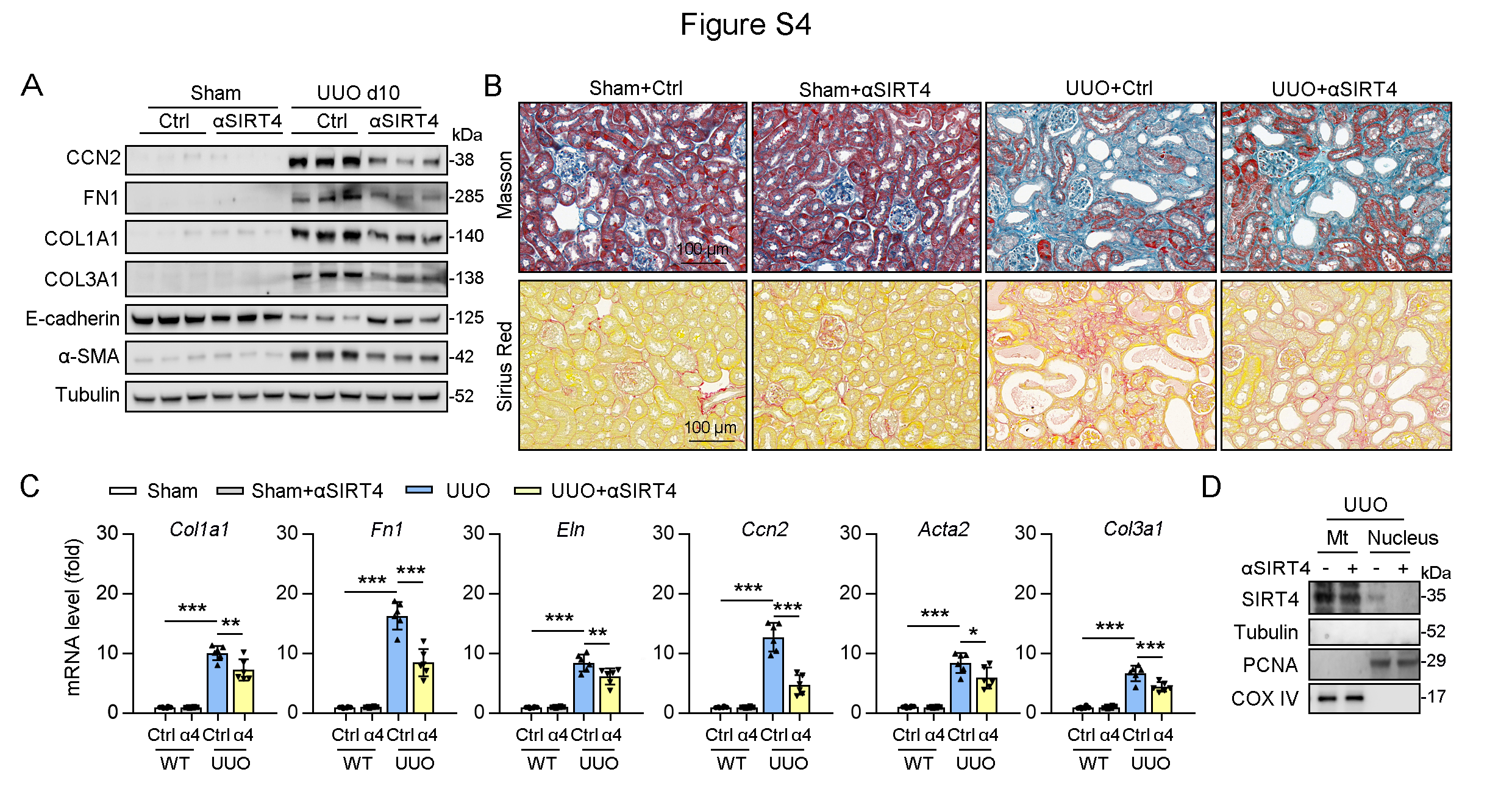

**Figure S4. Exosomes contain anti-SIRT4 antibody alleviated UUO-induced kidney fibrosis. (A to D)** After sham or UUO surgery, WT mice were treated with exosomes null or contain anti-SIRT4 for 10 days (n = 6 per group). A: Western blot analysis of the expression of CCN2, FN1, COL1A1, COL3A1, E-cadherin. α-SMA and Tubulin in the kidney from mice. B: Representative images of Masson’s trichrome staining and Sirius red staining in kidneys sections of mice (scale bar, 100 μm). C: The mRNA level of *Col1a1, Fn1, Eln, Ccn2, Acta2*, and *Col3a1* in the kidney of mice. D: Organelle separation experiment and immunoblot analysis detected the localization of SIRT4 in the kidneys of mice. For all panels, data are presented as mean ± SD. **P* < 0.05, ***P* < 0.01, ****P* < 0.001 by one-way ANOVA with Bonferroni correction test.

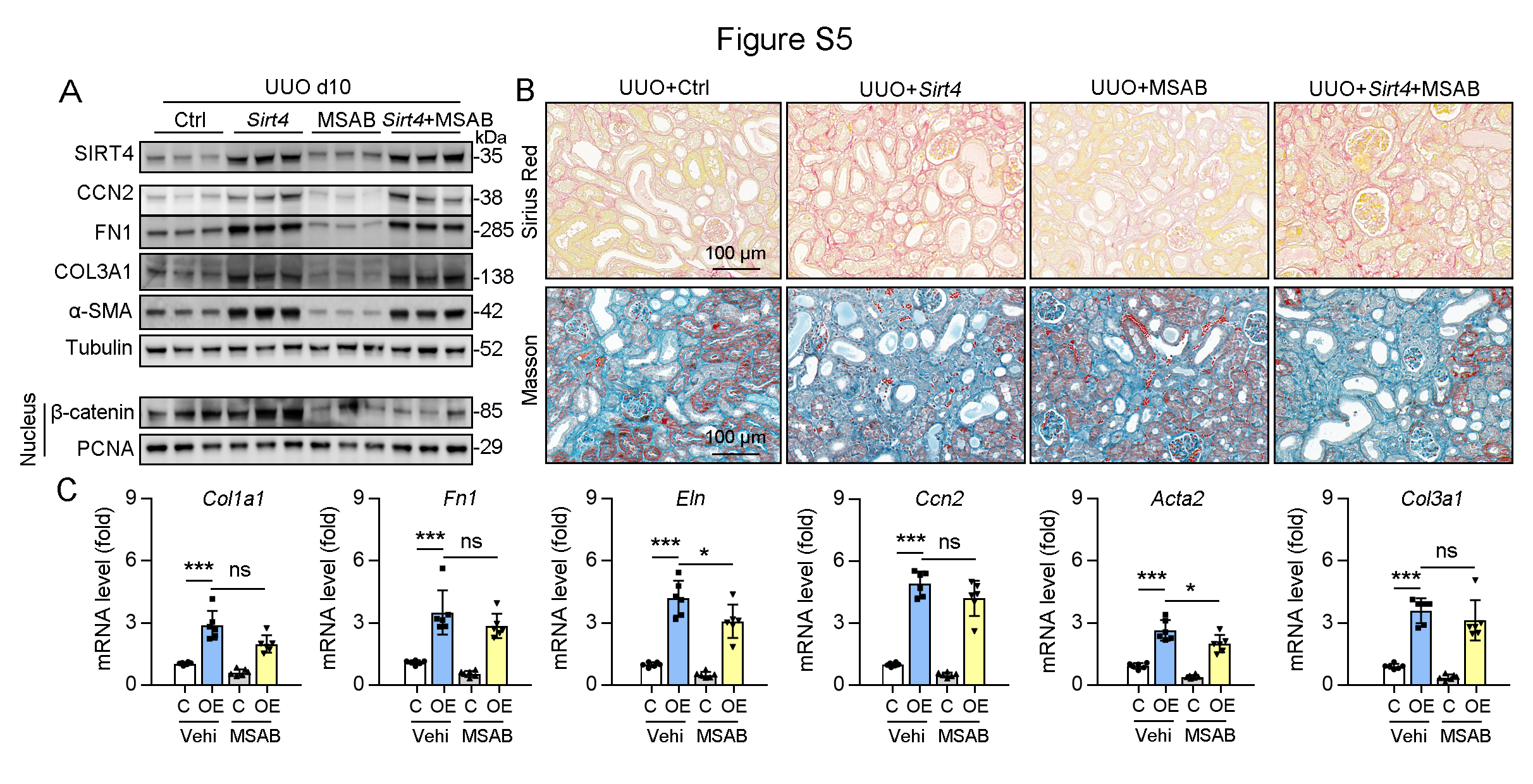

**Figure S5. Inhibition of nucleus accumulation of β-catenin failed to suppresses kidney fibrosis induced by SIRT4 overexpression in UUO mice. (A to C)** AAV9-*Ksp*-*Sirt4* or AAV-Ctrl was injected into kidneys of mice in situ at three independent points in situ. After 2-week transfection, the mice received UUO surgery. After UUO surgery, mice were treated with vehicle or MSAB for 10 days (n = 6 per group). A: Western blot analysis of the expression of SIRT4, CCN2, FN1, COL3A1, α-SMA, and Tubulin in the kidneys from mice. B: Representative images of Masson’s trichrome staining and Sirius red staining in kidney sections of mice (scale bar, 100 μm). C: The mRNA level of *Col1a1, Fn1, Eln, Ccn2, Acta2,* and *Col3a1* in the kidney of mice*.* For all panels, data are presented as mean ± SD. ns: not significant difference, **P* < 0.05, ***P* < 0.01, ****P* < 0.001 by one-way ANOVA with Bonferroni correction test.

**2. ESM Methods**

**2.1 Protein extraction and Western blot analysis**

Cytoplasmic or nuclear extracts were prepared from cells or kidneys using a cytoplasmic and nuclear protein isolation kit (Cat# 78833, Thermo Fisher Scientific). Briefly, the tube was vortexed vigorously in the highest setting for 15 s to fully suspend the cell pellet (or tissues were homogenized in PBS). The tubes were incubated on ice for 10 min. Next, ice-cold cytoplasmic extraction reagent II solution was added to the tubes. The tube was vortexed and incubated on ice. The tube was centrifuged for 5 min at maximum speed in a microcentrifuge (~16,000 × g), and the supernatant (cytoplasmic extract) was transferred to a tube. The insoluble (pellet) fraction, which contains nuclei, was suspended in ice-cold nuclear extraction reagent and then vortexed at the highest setting for 15 s. The sample was placed on ice and vortexed for 15 s every 10 min for 40 min. The tube was centrifuged at maximum speed (~16,000 × g) in a microcentrifuge for 10 min, and the supernatant (nuclear extract) fraction was transferred to a tube.

Mitochondria were extracted using a Mitochondria Isolation Kit (Sigma) following the manufacturer’s protocol. In brief, 2 ×10^7^ cells were pelleted by centrifuging the harvested cell suspension, and then mitochondria isolation reagent was added to the cell pellets. The cell resuspension was centrifuged at 700 × g for 10 min at 4°C, and then the supernatant was transferred to a new 1.5 ml tube and centrifuged at 12,000 × g for 15 min at 4°C. The supernatant (cytosolic fraction) was transferred to a new tube, and the pellet contained the isolated mitochondria. The pellet was further lysed to yield the final mitochondrial lysate. The extracted proteins were prepared for subsequent western blotting analysis.

For western blot analysis, 50 µg of lysate was loaded onto sodium dodecyl sulfate-polyacrylamide gel electrophoresis gels, and transferred onto polyvinylidene difluoride membranes (Millipore). Proteins were analyzed with their corresponding specific antibodies. Densitometry analysis was performed using Quantity One® Software and quantified relative to the loading control, Tubulin.

**2.2 Antibodies used in Western blottings**

| ANTIBODIES | SOURCE | IDENTIFIER | |
| --- | --- | --- | --- |
| SIRT4 (1：1000) | Thermo Fisher Scientific | | Cat# PA5-114377; RRID: AB_2787525 |
| SF3B1 (1:1000) | Thermo Fisher Scientific | | Cat# MA5-42938; RRID:AB_2912079 |
| KAT5 (1:1000) | Thermo Fisher Scientific | | Cat# PA5-34548; RRID:AB_2551900 |
| SF3B5 (1:1000) | Thermo Fisher Scientific | | Cat# A305-586A-T; RRID:AB_2782745 |
| SF3B3 (1:1000) | Abcam | | Cat# ab209402; RRID:AB_2910146 |
| CCN2 (1:1000) | Abcam | | Cat# ab6992; RRID:AB_305688 |
| FN1 (1:1000) | Abcam | | Cat# ab2413; RRID:AB_2262874 |
| α-SMA (1:1000) | Abcam | | Cat#ab7817; RRID:AB_262054 |
| U2AF2 (1:50/1: 250) | Abcam | | Cat# ab37530; RRID:AB_883336 |
| Flag (1:50/1:1000) | Abcam | | Cat# ab205606; RRID:AB_2916341 |
| Anti-acetyl Lysine (1:1000) | Abcam | | Cat# ab21623; RRID:AB_446436 |
| PCAF (1:1000) | Abcam | | Cat# ab96510; RRID:AB_10679933 |
| KAT8 (1:1000) | Abcam | | Cat# ab200660; RRID:AB_2891127 |
| COX IV (1:1000) | Abcam | | Cat# ab202554; RRID:AB_2861351 |
| KPNA2 (1:1000) | Abcam | | Cat# ab84440; RRID:AB_1860701 |
| p300 (1:1000) | Abcam | | Cat# ab275378; RRID:AB_2935873 |
| HAT1 (1:1000) | Abcam | | Cat# ab194296; RRID:AB_2801641 |
| BAK | Abcam | | Cat# ab104124; RRID:AB_10712355 |
| COL1A1 (1:1000) | Proteintech | | Cat# 67288-1-Ig; RRID:AB_2882554 |
| HA-Tag (1:50/1:1000) | Cell signaling Technology | | Cat# 3724; RRID:AB_1549585 |
| Poly/Mono-ADPRibose (1:1000) | Cell signaling Technology | | Cat# 83732; RRID:AB_2749858 |
| CBP (1:1000) | Cell Signaling Technology | | Cat# 7389; RRID:AB_2616020 |
| E-cadherin (1:1000) | Proteintech | | Cat# 20874-1-AP; AB_10697811 |
| Tubulin (1:1000) | Proteintech | | Cat# 11224-1-AP; RRID:AB_2210206 |
| PCNA (1:2000) | Novus | | Cat# NB500-106; RRID:AB_2252058 |
| PUF60 (1:1000) | Novus | | Cat# NBP1-49906; RRID:AB_10012096 |
| SF3B4 (1:1000) | Novus | | Cat# NBP1-92692; RRID:AB_11025842 |
| COL3A1 (1:1000) | Novus | | Cat# NB600-594; RRID:AB_10001330 |
| SF3B2 (1:1000) | Novus | | Cat# NBP1-92380; RRID:AB_11031649 |
| BAX (1:1000) | Novus | | Cat# NBP1-28566; RRID:AB_1852819 |
| ERK1/ERK2 (1:1000) | Novus | | Cat# AF1576; RRID:AB_354872 |
| p-ERK1/2 (1:1000) | Novus | | Cat# AF1018; RRID:AB_354539 |
| U2AF1（1:100/1:1000） | Novus | | Cat# NBP1-32515; RRID:AB_2288123 |

**2.3 Histological analysis**

Kidney tissue was embedded in paraffin and sliced into 5 μm thick serial sections using a paraffin slicer. For kidney histology, the paraffin sections were stained using Masson’s trichrome and Sirius red staining kit (Solarbio & Technology, Beijing, China).

**2.4 Quantitative real-time PCR**

Total RNAs were extracted by Trizol (Invitrogen) and then dissolved in an appropriate amount of RNase water. cDNA was obtained by a reverse transcription kit purchased from TransGen Biotech (Beijing, China). qPCR was performed using the ABI StepOnePlusTM Real-time PCR system (Applied Biosystem) with specific primers **(Table S2)**. The relative mRNA levels of target genes were analyzed using the 2^-ΔΔCT^ method. Tubulin was used as a housekeeping gene for analysis.

**2.5 RIME**

The procedure was performed as previously described. The hTECs (2 × 10^7^) were treated with DMSO or DMOG (1mM) for 8 h and cross-linked with 1% formaldehyde for 8min and quenched by 0.125M glycine. Cells were harvested and the nuclear fraction was extracted. After sonication, the cell lysate was immunoprecipitated with bead-prebound SIRT4 antibody (NB100-479, Novus Biologicals). Precipitated proteins in 30 μl of 100mM ammonium hydrogen carbonate were reduced in 2.5mM dithiothreitol at 60 °C for 1 h, then alkylated with 5mM iodoacetamide in the dark at room temperature for 15min. Proteins were digested with 20 ng/μl Trypsin/LysC (Promega) at 37 °C overnight. Peptides were desalted on Oasis HLB μ-elution plates (Waters), eluted with 65 % acetonitrile/0.1 % trifluoroacetic acid, dried by vacuum centrifugation, and analyzed by liquid chromatography/tandem mass spectrometry using a nano-Easy LC 1000 interfaced with an Orbitrap Fusion Lumos Tribrid Mass Spectrometer (Thermo Fisher Scientific). Tandem mass spectra were extracted by Proteome Discoverer version 2.3 (Thermo Fisher Scientific) and searched against the SwissProt_Full_Synchronized_2018_08 database using Mascot version 2.6.2 (Matrix Science). Mascot “.dat” files were compiled in Scaffold version 3 (Proteome Software) to validate MS/MS-based peptide and protein identifications. Peptide identifications were accepted if false discovery rate (FDR) was less than 1%, based on a concatenated decoy database search by the Peptide Prophet algorithm with Scaffold delta-mass correction.

**2.6 Co-immunoprecipitation (co-IP)**

After treatment, the cells were lysed in an ice-cold co-IP buffer containing 20 mM Tris-HCl (pH 8.0), 100 mM NaCl, 1 mM EDTA, and 0.5% NP-40, supplemented with a protease inhibitor cocktail (Roche, 04693132001), for 30 min. The cell homogenates were then centrifuged at 13,000g for 15 min, and the resulting supernatant was incubated overnight at 4 °C on a shaker with primary antibodies (anti-Flag, anti- SIRT4, anti-HA, anti-U2AF2 or anti-IgG). To ensure complete saturation of the primary antibodies, sufficient cell lysates were cultured and collected for IP. The mixture of antibodies and proteins was subsequently incubated with protein A/G-agarose beads (Thermo Fisher Scientific, Cat#: 78610) at 4 °C for 3 h. The beads were washed 5-6 times with cold IP buffer and resuspended in loading buffer. The cell lysates and immunoprecipitates were denatured in loading buffer at 95 °C for 5 min, and western blotting analysis was performed.

**2.7 Protein-protein docking simulation**

Visual protein-protein docking of SIRT4 and U2AF2 or ERK2 was performed online using HDOCK (http://hdock.phys.hust.edu.cn/).

**2.8 Analysis of the conservation of lysine 413 in U2AF2**

A gene panel of the NCBI database (https://www.ncbi.nlm.nih.gov/gene/) was used to download amino acid sequences of proteins from multiple species and subsequently compare sequence conservation between species using the UGENE software (version 39; the software can be downloaded from the following website: http://ugene.net/).

**2.9 Bioinformatics methods predict acetylase of Lys413 in U2AF2**

A Group-based Prediction System (GPS) online tool (http://pail.biocuckoo.org) was used to predict Lys413 in U2AF2 modified by histone acetyltransferases, with high score values suggesting better results. The amino acid sequence of the protein in FASTA format was downloaded from UniProt and uploaded to the GPS website.

**2.10** **Luciferase report assay for TGF-β1**

Specific primers for the 5’-ACTCTCGAGCAGTGTTCCCACCCTGACAC-3’ and for 5’-ACTAAGCTTGGTTGGCACTGCGGGCGGAG-3’ were designed to amplify the DNA (-835/+214) in the CTGF promoter region of the human genome through the polymerase chain reaction (PCR) experiment. The sequence of the human CTGF promoter was cloned into the pGL3.0-Basic vector (LMAI Bio; Cat# LM-1554) through molecular cloning. Positive plasmids were screened and identified by methods such as monoclonal colony PCR, plasmid double enzyme digestion identification, and sequencing. Then, using Lipofectamine 3000 transfection reagent, pGL3.0-Basic-CTGF and Renilla luciferase were transfected into HEK293T. Firefly and Renilla luciferase activities were then measured by a dual luciferase reporter gene system (Promega, Madison, WI, USA)

**2.11 Cell culture, transfection**

The human proximal tubular epithelial cell line (HK2 cells) (Cat# CRL-2190) were purchased from the American Type Culture Collection (Manassas, VA, USA). Cells were cultured in Minimum Essential Medium (Thermo Fisher Scientific, Shanghai, China; Cat# 10373017). Mouse podocytes (MPC) (Cat# BNCC342021) and mouse renal glomerular endothelial cells (GEC) (Cat# BNCC360313) were purchased from BeNa Culture Collection (Beijing, China), cultured in DMEM-H complete medium (BeNa Culture Collection, Cat# BNCC338068). Mouse renal fibroblasts (MF) (Procell Life Science&Technology, Wuhan, china; Cat# CP-M069) were cultured in mouse kidney fibroblast complete culture medium (Procell, Cat#CM-M069). All kinds of media were supplemented with 10% FBS (Gibco, Grand Island, NY), 100 U·mL−1 of penicillin (Gibco), and 100 mg•mL−1 of streptomycin (Gibco). All cells were cultured in a 37 °C incubator containing 5% CO2 and 95% air.Human embryonic kidney 293 cells (HEK293) were purchased from ATCC and maintained in HyClone Dulbecco’s Modified Eagle Medium (SH30022, Cytiva) with 10 % FBS, 1 % glutamine, and 1 % penicillin/streptomycin solution in a humidified incubator supplemented with 95 % air/5 % CO2 at 37 °C. Cells were regularly checked for mycoplasma in a standardized manner, by a qPCR test, performed under ISO17025 accreditation to ensure work was conducted in mycoplasma-negative cells.

Adenovirus particles used in the article were provided by GeneChem Company (Shanghai, China). Adenovirus was used to transfect HK2 cells or TECs for 6 h (adenovirus particles containing empty vectors were used as controls), and then replace the fresh medium and continue to culture. The commercial siRNA Ctrl (Cat# sc-37007, Santa Cruz), siRNA *Hat1* (Cat# sc-145898, Santa Cruz), siRNA *Sirt4* (Cat# sc-63025, Santa Cruz), siRNA *Bax* (Cat# sc-29213, Santa Cruz), and siRNA *Bak* (Cat# sc-29785, Santa Cruz) were transfected using Lipofectamine 3000 (Invitrogen, Carlsbad, California, USA) according to the manufacturer’s instructions.

**2.12 Isolation of primary renal tubular epithelial cells (TECs)**

Primary mouse renal tubular epithelial cells were isolated using an established protocol we previously published^1^. Briefly, the cortex of the kidneys was carefully dissected and chopped into small pieces. Then, 1 mg/ml of collagenase solution was applied and incubated at 37 °C for 30 min with gentle agitation. The digestion was terminated by FBS and then filtered sequentially. Fragmented tubules were collected and maintained in a renal epithelial cell basal medium using a growth kit. The medium was changed for the first time, after 72 h. The purity of the primary mouse renal tubular epithelial cells was confirmed by immunofluorescence staining of SGLT1 (Novus, NBP2-20338). Cells at passages 2-5 were used for the experiments.

**2.13 Immunofluorescent staining of cells**

After treatment, cells was fixed for 15 min using fresh, methanol-free 4% formaldehyde, and then, rinsed thrice with PBS for 5 min each. After blocking with goat serum for 30 min, the cells were incubated with primary antibody against SIRT4 (Thermo Fisher Scientific, Cat# PA5-114377) at 4 °C overnight. Alexa Fluor®488 goat antibodies against murine IgG (Invitrogen, Shanghai, China; Cat# A-11078, 1:400) were included as secondary antibodies. As negative controls, the primary antibodies were exchanged for nonimmune serum from the same species. The samples were counterstained with DAPI for 15 min. The sections were sealed with a cover glass, and the specimens were examined using the appropriate excitation wavelength. Images were captured and processed with a Laica microscope (Wetzlar, Germany).

**2.14 Immunohistochemical (IHC) staining**

Paraffin-embedded mouse kidney samples were sliced into 4 μm-thick sections and subjected to IHC staining using a Rabbit two-step detection kit (Rabbit enhanced polymer detection system, ZSGB Bio, Beijing, China). Briefly, mouse kidney sections were deparaffinized, followed by antigen retrieval, treatment with peroxidase block, and incubation with rabbit anti-SIRT4 (1:100, Thermo Fisher Scientific, Cat# PA5-114377) primary antibody overnight at 4°C. Tissue sections were then washed and incubated with the reaction-enhancing solution at room temperature for 20 min. After washing three times with phosphate buffer saline (PBS), tissue sections were treated with enhanced enzyme-labeled goat anti-rabbit IgG polymerase for 20 min and then developed with 3,3' Diaminobenzidine solution, counterstained with hematoxylin, and mounted with mounting medium. The kidney frozen slices from patients with minimal change disease and DKD were fixed with 4 % paraformaldehyde for 20 min, incubated with anti-SIRT4 primary antibody overnight at 4 °C, and subjected to immunohistochemistry staining as described above. The stained area was measured using ImageJ software.

**2.15 RNA immunoprecipitation (RIP)**

RIP is a powerful method to investigate the interactions between RNA and RNA binding proteins in vivo by immunoprecipitating RNA-protein complexes using specific antibodies against RNA binding proteins. An example of the RIP protocol is as follows: Covalently conjugate protein-specific antibody to the beads. Wash extensively with lysis buffer. Collect cell lysate and incubate with antibody conjugated beads allowing RNA-protein complexes formation. Use normal IgG as negative control. Incubate the lysate for 2-4 hours at 4°C with gentle rotation. Wash beads stringently with ice-cold lysis buffer containing RNase inhibitors, protease inhibitors and nonionic detergents. Perform 4-6 quick washes. Elute RNA-protein complexes from beads, generally with 1% SDS buffer heated to 95°C for 5min. Extract RNA with phenol/chloroform. Isolate RNA for downstream applications like RT-qPCR or RNA sequencing.

**2.16 The stability and intron retention index of CCN2 mRNA**

To detect the CCN2 mRNA stability, the cells were treated with Actinomycin D (2 mg/mL) for 0 h, 1 h, 2 h and 3 h. The cDNA from different time point was diluted in nuclease-free water for the same times, and mRNA was quantified by qPCR. The fold of changing CCN2 level was normalized to the initial expression level of CCN2 (0 h). Intron Retention index = (mRNA level of normal splicing isoform/total CCN2 mRNA level)/ (mRNA level of abnormal splicing isoform/total CCN2 mRNA level). The CCN2 mRNA stability qPCR primers are as listed in Table S3.

**2.17 Loading of Exosomes**

The approaches for αSIRT4 incorporation into exosomes were executed as mentioned before^3^. Naive exosomes released from HEK293T were diluted in PBS to a concentration 0.15 mg/mL of total protein, then αSIRT4 solution in PBS (0.5 mg/mL) was added to 250 μl of exosomes to the final concentration 0.1 mg/mL total protein, and incubated at RT for 18 hours. In case of a saponin treatment, a mixture of αSIRT4 and exosomes was supplemented with 0.2% saponin and placed on shaker for 20 min at room temperature.

**3. Table**

**Table S1. Clinical characteristics of CKD in biopsy samples, Related to Figure 1**

| Parameters | Mild CKD (n=4) | Severe CKD (n=4) | *P* value |
| --- | --- | --- | --- |
| Sex | Male:Female=3:1 | Male:Female=2:2 | NS |
| Age | 36.3±7.8 | 47.5±17.5 | NS |
| FBG (mmol/L) | 6.57±1.69 | 5.82±0.81 | NS |
| Proteinuria (g/24h) | 2.49±2.01 | 10.96±7.95 | 0.042 |
| eGFR (ml/min/1.73m^2^) | 84.42±27.49 | 38.875±12.94 | 0.012 |
| Duration of disease (y) | 4±2.3 | 5±4.8 | NS |

Data are calculated by One-way ANOVA and presented as mean ± SEM. The number of patients in each group was as indicated. NS: not significant difference, y: year.

**Table S2. Primers used in qPCR**

| Gene | Forward Primer (5’→3’) | Reverse Primer (5’→3’) |
| --- | --- | --- |
| m*Col1a1* (ID: 12842) | *GCTCCTCTTAGGGGCCACT* | *CCACGTCTCACCATTGGGG* |
| m*Col3a1* (ID: 12825) | *CTGTAACATGGAAACTGGGGAAA* | *CCATAGCTGAACTGAAAACCACC* |
| m*Fn1* (ID: 14268) | *ATGTGGACCCCTCCTGATAGT* | *GCCCAGTGATTTCAGCAAAGG* |
| m*Eln* (ID: 13717) | *TTGCTGATCCTCTTGCTCAAC* | *GCCCCTGGATAATAGACTCCAC* |
| m*Ccn2* (ID: 14219) | *GGGCCTCTTCTGCGATTTC* | *ATCCAGGCAAGTGCATTGGTA* |
| m*Acta2* (ID: 11475) | *CCCAACTGGGACCACATGG* | *TACATGCGGGGGACATTGAAG* |
| m*Ngal* (ID: 16819) | *TGGCCCTGAGTGTCATGTG* | *CTCTTGTAGCTCATAGATGGTGC* |
| m*Kim-1* (ID: 171283) | *GTTAAACCAGAGATTCCCACACG* | *TCTCATGGGGACAAAATGTAGTG* |
| m*β-actin* (ID:11461) | *GGCTGTATTCCCCTCCATCG* | *CCAGTTGGTAACAATGCCATGT* |

**Table S3. Primers used in IR Index**

| *CCN2* pre-mRNA | Forward Primer (5’→3’) | Reverse Primer (5’→3’) |
| --- | --- | --- |
| In 2+ex 3 | *GCCTATTCTGTCACTTCGGC* | *CACTCCTCGCAGCATTTCC* |
| In 3+ex 4 | *TCAGGGTCGTGATTCTCTCC* | *TCTCTTCCAGGTCAGCTTCG* |
